# FORK-seq: replication landscape of the *Saccharomyces cerevisiae* genome by nanopore sequencing

**DOI:** 10.1101/2020.04.09.033720

**Authors:** Magali Hennion, Jean-Michel Arbona, Laurent Lacroix, Corinne Cruaud, Bertrand Theulot, Benoît Le Tallec, Florence Proux, Xia Wu, Elizaveta Novikova, Stefan Engelen, Arnaud Lemainque, Benjamin Audit, Olivier Hyrien

## Abstract

Genome replication mapping methods profile cell populations, masking cell-to-cell heterogeneity. Here, we describe FORK-seq, a nanopore sequencing method to map replication of single DNA molecules at 200 nucleotide resolution. By quantifying BrdU incorporation along pulse-chased replication intermediates from *Saccharomyces cerevisiae*, we orient 58,651 replication tracks reproducing population-based replication directionality profiles and map 4,964 and 4,485 individual initiation and termination events, respectively. Although most events cluster at known origins and fork merging zones, 9% and 18% of initiation and termination events, respectively, occur at many locations previously missed. Thus, FORK-seq reveals the full extent of cell-to-cell heterogeneity in DNA replication.

## INTRODUCTION

Eukaryotic DNA replication initiates at multiple replication origins and terminates wherever converging replication forks happen to meet. Understanding this process is essential as replication perturbations can threaten genome stability. DNA microarrays and massive DNA sequencing techniques have triggered an explosion of genome-wide replication mapping studies in the last decade. However, these cell population-based methods only provide an average profile of DNA replication where cell-to-cell heterogeneity is masked. A high-throughput single-molecule method is required to reveal this heterogeneity.

DNA combing, the modern, fluorographic version of DNA fibre autoradiography, relies on antibody detection of nucleotide analogs incorporated during replication, combined with *in situ* hybridization of stretched DNA molecules with DNA probes, to reveal the replication pattern of single copies of a locus of interest [1, 2]. While this method has produced valuable information at selected loci, it is technically difficult and very low throughput. Recent improvements were achieved by using fluorescent dNTPs to directly label newly replicated DNA and barcode total DNA at nicking endonuclease sites [3] followed by DNA stretching in nanochannel arrays developed for automated genome assembly [4, 5]. Nevertheless, DNA stretching inhomogeneities, optical resolution and image processing challenges still restrain the power of optical replication mapping.

MinION nanopore sequencing (Oxford Nanopore Technologies, ONT) has the potential to bypass these limitations. This novel technology produces long (up to 2.3 Mb [6]) reads of native single-stranded DNA molecules by measuring changes in ionic current across a nanopore through which DNA is translocated. Importantly, the nanopore sequencer can detect base modifications on long RNA and DNA molecules [7, 8]; for review [9]. In particular, we and others reported the detection of bromodeoxyuridine (BrdU) or other thymidine analogs incorporated during DNA replication in yeast or mouse DNA [10, 11, 12]. Using *S. cerevisiae* cells synchronously progressing through S phase in conditions of limiting BrdU concentrations, Müller *et al* [11] detected gradients of BrdU incorporation along single DNA molecules at 2 kb resolution. Importantly, most but not all peaks of BrdU incorporation mapped near known origins, suggesting the existence of dispersed origins too rarely used to be detected by cell population methods.

Here, by precisely quantifying BrdU incorporation at 200 nucleotide-resolution, we detected, mapped and oriented short BrdU incorporation tracks synthesized during a brief (4 min) pulse followed by a chase in asynchronously growing *S. cerevisiae* cells. We named this method FORK-seq. Analysis of 58,651 oriented tracks not only reproduced replication fork directionality (RFD) profiles independently obtained by sequencing of Okazaki fragments (OK-seq) but also identified 4,964 and 4,485 individual initiation and termination events, respectively. The majority of initiation events formed clusters that coincided with known origins. However, 9% of initiation events mapped away from known origins, at mainly dispersed locations that typically lacked the consensus sequence and origin recognition complex (ORC) and Mcm2-7 binding peaks found at known origins. Termination events were more dispersive than previously recognized, as a large number of them occurred outside fork merging zones previously detected in cell population studies [13, 14]. These results further illustrate the power of nanopore sequencing for mapping genome replication by single molecule analysis, providing information unreachable by cell-population methods. They support a model in which replication of eukaryotic chromosomes combines clustered initiation at efficient sites associated with specific DNA sequences, with dispersed initiation at inefficient sites that lack sequence specificity and inefficiently recruit ORC and Mcm2-7.

## RESULTS

### BrdU produces a distinct nanopore electrical signal

The ONT MinION instrument measures changes in electrical current as a single DNA strand is translocated through a protein pore to reveal DNA sequence. Several consecutive nucleobases in the narrowest region of the pore can influence the ionic current. Translating a sequence of current values into a DNA sequence is therefore a non-trivial task typically solved using hidden Markov models [15, 16] or recurrent neural networks [17, 18]; for review [9]. Importantly, such approaches can discriminate methylated and hydroxymethylated from unmodified cytosines [7, 8], suggesting that detection of modified nucleobases incorporated in newly replicated DNA should be feasible. To measure the effect of BrdU incorporation on the current signal, we generated control or BrdU-hemisubstituted DNA duplexes by primer extension of linearized plasmid DNA in the optional presence of dTTP or BrdUTP, followed by exonuclease degradation of the non-template strand (Fig. 1a). Bioanalyzer and Qubit analyses (Fig. 1b) revealed a high yield of primer extension and an electrophoretic shift of BrdU-substituted DNA. A small amount of duplex DNA was observed in the absence of dTTP and BrdUTP, likely due to partial plasmid renaturation before exonuclease degradation.

**Figure 1:**
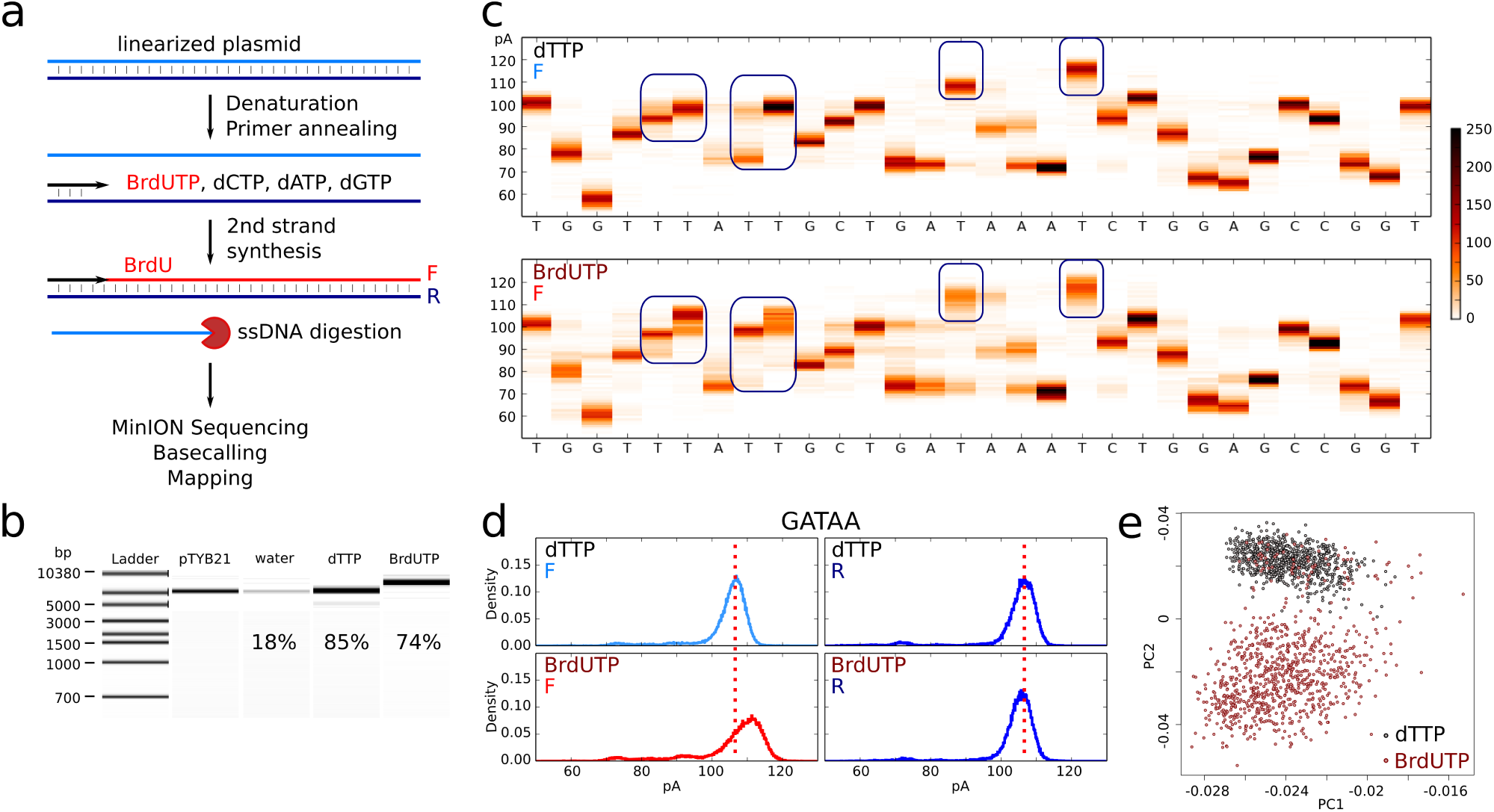
Effect of BrdU incorporation into DNA on nanopore sequencing current signal. a. Scheme of sample preparation. F, forward strand; R, reverse strand. b. Bioanalyzer size control of the samples, with Qubit yield indicated. pTYB21, linearized plasmid; water, primer extension in the absence of dTTP and BrdUTP; dTTP, primer extension using canonical dNTPs; BrdUTP, primer extension using BrdUTP instead of dTTP. c. Example of a 30 bp sequence of the forward (F) strand (positions 1000-1029) with current distribution of 500 reads at each position. Upper panel: sample obtained using canonical dNTPs. Lower panel: dTTP was replaced by BrdUTP. Blue rectangles highlight some current shifts due to the presence of BrdU. BrdU did not induce a current shift at all thymidine sites. d. Current distribution for the ‘GATAA’ pentamer for the dTTP (top) and the BrdUTP (bottom) sample on the forward (F, modified strand, left) and the reverse (R, native strand, right) strands. e. Principal component analysis using as inputs 1kb long current value sequences (positions 100-1100 on the reference plasmid sequence) from 1000 reads for dTTP (black) and BrdUTP (brown) samples (F strand). The first two components are represented. Only “pass” reads were used in c,d,e.

The primer extension products were sequenced using the MinION (R9 chemistry) and the “2D” protocol where the two strands of a DNA duplex are consecutively read thanks to a hairpin adapter. Sequencing information is summarized in Additional file 1: Table T1. The raw data were basecalled with Metrichor (ONT) and the sequences of the two complementary strands were independently aligned to the plasmid using BWA MEM [19] with parameters adapted to the sequencing error rate (see Materials and methods, Additional file 2: Fig. S1). Since Metrichor was devised to detect canonical bases, the presence of BrdU may affect basecalling and mapping. Indeed, only 49% of the BrdUTP vs. 60% of the dTTP reads could be mapped (Additional file 1: Table T2). In the 2D protocol, Metrichor classifies the reads into “pass” and “fail” based on the presence of a second strand read and its complementarity to the first strand. While the fraction of 2D reads was similar for both samples (57% vs. 59%), the percentage of “pass” reads was lower for BrdUTP (13%) than for dTTP (18%), indicating lower read complementarity. Importantly, 99% of the “pass” reads were successfully mapped for both samples (Additional file 2: Fig. S1b and Additional file 1: Table T2). However, the parental (reverse, R) and newly replicated (forward, F) strand miscall rates were similar for dTTP (7% vs. 7%) but not for BrdUTP (7% vs. 11%, respectively, Additional file 2: Fig. S1c), confirming that BrdU affects the current in the pore. In conclusion, current alterations due to BrdU reduce Metrichor basecalling accuracy but to an extent that does not strongly affect read alignment to the reference sequence.

Python scripts were then developed to realign current steps to the reference DNA sequence. To allow comparisons between multiple experiments, we normalized each profile by subtracting its mean current intensity. As shown on Fig. 1 (c and d), thymidine (T) was associated in most cases with the highest current values in native DNA, and its substitution by BrdU further increased the current at these sites but not at other bases. This probably explains why BrdU did not cause systematic basecalling errors at T sites (Additional file 2: Fig. S1c). We extracted for each pentamer the difference in median current value between BrdUTP and dTTP samples (Additional file 2: Fig. S1d). Almost all F-strand pentamers with a T in their middle displayed a positive current shift in the presence of BrdU, with 60% showing a shift > +3 pA. This effect was not seen when the T was at another position and most pentamers lacking this central T showed a small negative shift in the BrdU sample, due to subtraction of a higher mean current than in the dTTP sample during normalisation. Importantly, only the BrdU-substituted (F) strand was perturbed, while the complementary native (R) strand gave an identical signal to the thymidine control (Fig. 1d, Additional file 2: Fig. S1c,d). A principal component analysis of current values aligned on a 1 kb reference fragment showed a clear separation of BrdU-substituted and control fragments (1000 of each, Fig. 1e). The small number of BrdUTP sample reads that clustered with control reads most likely originated from unsubstituted DNA duplexes formed by partial renaturation of the native plasmid strands (visible in Fig. 1b, ‘water’ lane). Overall, these results indicate that BrdU detectably alters the nanopore electric signal, and that its signature should be identifiable using appropriate algorithms. Since our preliminary report [10], other laboratories also reported detection of BrdU [11] or multiple thymidine analogs [12] in synthetic DNA templates.

### Detecting BrdU incorporation in genomic DNA

To determine whether BrdU can also be detected in labelled genomic DNA, we used the MCM869 yeast strain, which has been genetically modified to depend on exogenous thymidine for growth [20]. Adding BrdU to culture medium allowed cell growth, albeit at a slower rate than thymidine (Additional file 2: Fig. S2). Cells stopped proliferating after 3 cycles. We therefore expected ∼87.5% of genomic DNA strands to contain only BrdU, with the remaining ∼12.5% corresponding to the starting parental strands containing thymidine. Accordingly, the total BrdU content of this sample measured by mass spectrometry was 91% (Additional file 1: Table T3). We also prepared a thymidine-only control and 3 samples of intermediate BrdU content (see Materials and methods), the total BrdU contents of which were 0%, 9%, 38% and 69%, respectively, as determined by mass spectrometry (Additional file 1: Table T3). The five samples were sequenced on the MinION using R9.4.1 pore version. Replicates of the unsubstituted and highly substituted samples were also sequenced using the R9.5 version (discontinued). Reads were basecalled by Albacore (ONT) and aligned on the S288C reference genome using Tombo (ONT) which uses minimap2. We obtained 93k-600k reads per sample, 54 to 88% of which being mapped (Additional file 1: Table T3). Tombo resquiggle command also realigned current intensities on the reference genome. As seen for the *in vitro* template, the current was positively shifted by the presence of BrdU (Additional file 2: Fig. S3a).

To estimate BrdU incorporation along DNA molecules, we developed two independent methods based on normalized current shifts between successive current plateaus (Fig. 2a) that we integrated in a software called RepNano (Materials and methods, [21]). We first implemented a machine learning approach to convert a segment of 96 consecutive current shifts and their aligned reference sequence into the proportion of T sites that incorporated a BrdU in the segment. We adopted a convolutional neural network (CNN) architecture with 3 convolutional layers (Additional file 2: Fig. S4). The learning was performed using 4000 reads from each sample described above (28,000 reads in total). Training reads were presented as sets of non-overlapping 96 bp segments associated with their respective BrdU content. We then used the trained network to estimate BrdU content in the remaining reads. To reduce the noise level, we computed BrdU content in overlapping 96 bp windows shifted by 10 bases and averaged the results at each base. We plotted the distributions of measured BrdU content by 500 bp windows (Additional file 2: Fig. S3b). The sample containing 0% of BrdU showed a single peak of low BrdU content while the sample with 91% showed two peaks of high and low BrdU content, corresponding to replicated and parental DNA, respectively. Therefore, the CNN resolved substituted from non-substituted strands according to their BrdU content. Overall, the mean BrdU content estimated from sequence reads was in good agreement with the mass spectrometry measurement for every sample (Additional file 2: Fig. S3c).

**Figure 2:**
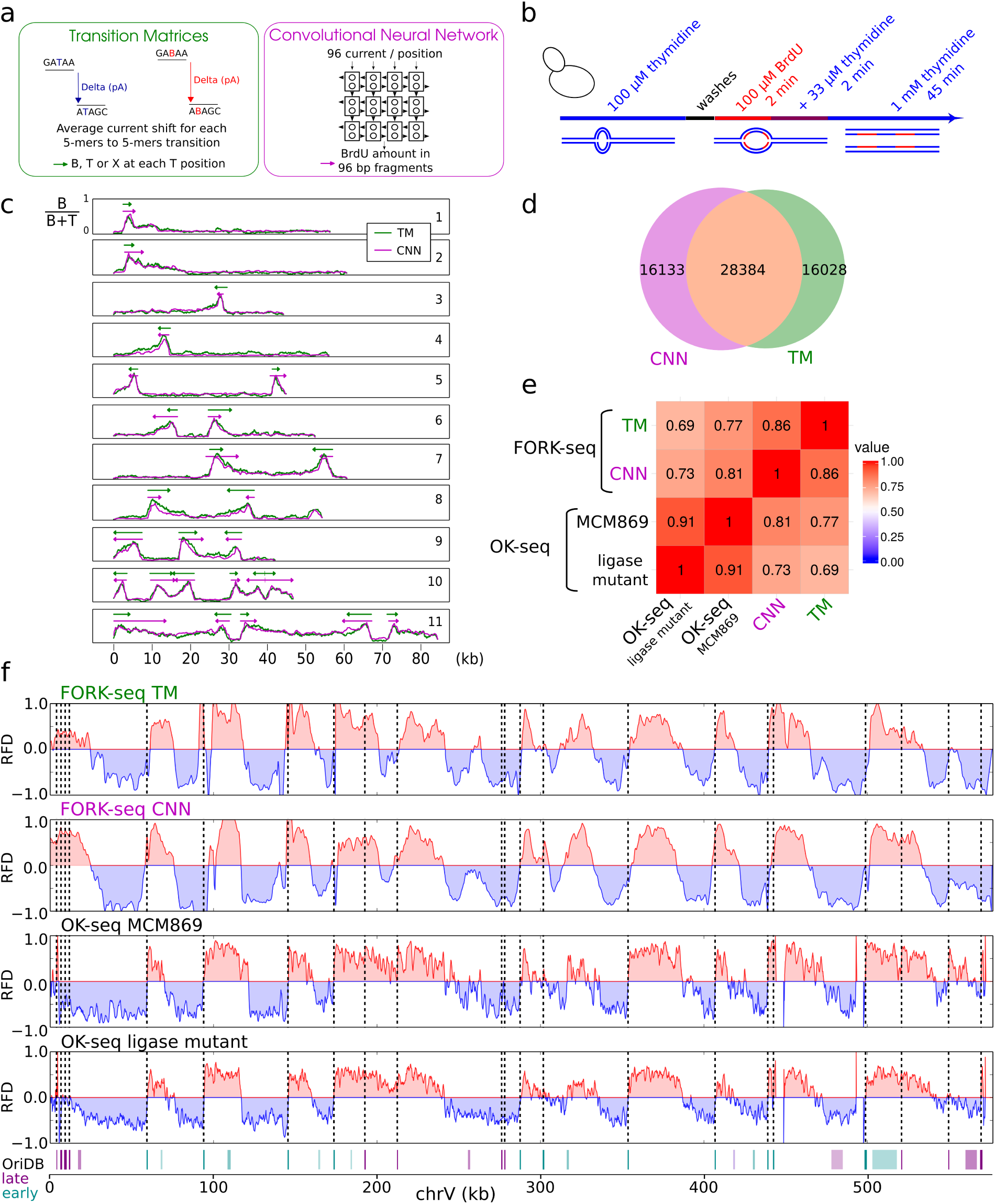
Detection of individual replication forks. a. BrdU content estimation methods. b. Scheme of sample preparation. c. Exemplary BrdU content profiles (B/(B+T) ratio; B, BrdU, T, thymidine) along nanopore reads using CNN (pink) or TM (green). Shown are examples for rightward(1,2) and leftward-moving (3,4) forks, initiation (5,6) and termination (7,8) events, and multi-replicon patterns (9-11). d. Venn diagram of the overlap between the CNN- and TM-detected forks. e. Spearman correlation coefficients between four RFD profiles of the whole genome at 1 kb smoothing obtained by the indicated methods. Top to bottom: FORK-seq TM and CNN, sequencing of EdU-labelled Okazaki fragments from MCM869 (OK-seq) and of accumulated Okazaki fragments from a ligase mutant [14]. f. RFD profiles from the four different methods for chromosome V. Bottom, known origins from OriDB [23] (violet, late, cyan, early; color intensity reflects OriDB classification: bright, confirmed, medium, likely, light, dubious). Vertical dotted line, confirmed origins.

The second method implemented in RepNano to estimate BrdU content is based on the calculation of the average current shift for every pentamer-to-pentamer transition in either thymidine or fully substituted BrdU reads (Fig. 2a). This produced two transition matrices (TM) that were used to predict, for every read and at each T position in the reference, if the measured current was most probably coming from a T or a B. Overall, distinction between BrdU and thymidine could be clearly made at one fifth of the T positions on the reference (Additional file 2: Fig. S3d). The remaining indeterminate positions were labelled X and treated as non-assigned bases. We then estimated BrdU content in the 0% and 91% samples (Additional file 2: Fig. S3b). The resulting distributions were more spread than with the CNN, but otherwise similar. As for CNN, there was a good correlation between TM estimates of the mean BrdU contents and mass spectrometry measurements for every sample, although both methods tended to underestimate BrdU content (Additional file 2: Fig. S3c). In conclusion, these results suggest that both CNN and TM methods can measure BrdU abundance along sequence reads.

### Detection of individual replication forks

We then tested whether our BrdU detection procedures were sufficiently precise to monitor replication fork progression during brief pulse-chase experiments. Exponentially growing MCM869 cells were pulsed with 100 μM BrdU for 2 min, added with 33 μM thymidine for 2 min and chased with 1 mM thymidine for 45 min (Fig. 2b). The low thymidine pulse preceding the chase was designed to obtain an intermediate incorporation of BrdU, allowing to determine the direction of fork progression. Genomic DNA was extracted and sequenced on the MinION (Additional file 1: Table T4) and BrdU content profiles were computed using CNN and TM. Both methods gave highly similar signals, reciprocally validating each other (Fig. 2c). The signals did not display plateaus, as it could have been expected if external nucleoside transport and phosphorylation were instantaneous. Instead, they showed a continuous variation of BrdU content, with a steep increase from ∼0% to 60-80% BrdU and a shallower decrease to ∼10% BrdU. We interpret the steep segments to reflect a rapid increase in the intracellular BrdUTP/dTTP ratio during the BrdU pulse, and the shallow segments to reflect a progressive decrease in this ratio during the final chase. Therefore signal asymmetry may immediately reveal the direction of replication fork progression.

To test this interpretation, we developed Python scripts to automatically detect and orient BrdU tracks, and compared the resulting replication fork directionality (RFD) profiles with independent RFD profiles generated by sequencing purified Okazaki fragments [14, 22]. Two biological replicates of the pulse-chase experiment were sequenced three times each (MinION, R9.4.1) to yield 1.4 million reads (Additional file 1: Table T4), 2.3% of which showed replication signals. A geometric method that approximates the signal by piecewise linear segments was used to detect replication tracks and infer orientation (Materials and methods), yielding 44,517 CNN and 44,412 TM oriented tracks, respectively, with 28,384 (64%) shared tracks (Fig. 2d). The incomplete overlap of the two track populations was due to the stringent thresholds used for detection and orientation. Of note, when run on native DNA with no BrdU, our algorithm output no fork, showing that the false positive rate was null. Replicate RFD profiles assembled from oriented BrdU tracks were well correlated to each other for each method (Spearman correlation 0.66 for TM and 0.75 for CNN for RFD computed from 1 kb smoothed data). The reads from the two replicates were therefore merged for further analyses. CNN- and TM-based RFD profiles were compared to each other and to two independent RFD profiles obtained by (i) sequencing overaccumulated Okazaki fragments from a ligase mutant [14] and (ii) sequencing purified, ethynyl-deoxyuridine (EdU)-labelled Okazaki fragments (OK-seq) [22] from the MCM869 strain (Fig. 2e,f). The four RFD profiles were strikingly similar (Spearman pairwise correlations 0.69 to 0.91, Fig. 2e). The correlation between the TM and CNN profiles (0.86) was almost as high as between the two OK-seq profiles (0.91). As expected, the TM and CNN profiles were better correlated to the MCM869 OK-seq profile (0.77 and 0.81, respectively) than to the ligase-deficient profile (0.69 and 0.73, respectively, Fig. 2e). As previously shown [14], initiation regions detected as ascending RFD segments coincided with the position of known yeast origins (from OriDB [23], Fig. 2f). These results demonstrate that our BrdU detection methods (CNN and TM) and track orientation procedures are robust and precise and that nanopore sequencing of pulse-chase labelled BrdU replication tracks, hereafter referred to as FORK-seq, is a valid alternative to OK-seq to produce genome-wide RFD profiles.

For comparison purposes, we attempted to use the D-NAscent software [11] to detect BrdU tracks on the same reads as CNN and TM. Although significant BrdU incorporation was detected at similar positions to CNN and TM, the signal was weaker (Additional file 2: Fig. S5). Using the same algorithm as for CNN or TM signals, most D-NAscent tracks could be neither detected nor oriented even after trying to optimize the detection parameters (Additional file 2: Fig. S5). The RFD profile assembled from the oriented D-NAscent tracks we obtained was of very poor quality, showing very low correlation coefficients to the other RFD profiles: FORK-seq TM, 0.15; FORK-seq CNN, 0.13; OK-seq MCM869, 0.12; OK-seq ligase mutant, 0.11. Therefore, in our hands, RepNano outperforms the DNAscent tools for precise measurements of BrdU incorporation and RFD profile establishment.

### FORK-seq precision

The first source of error in BrdU track detection comes from estimating BrdU incorporation along reads. Thus we looked at the average and standard deviation of CNN signals at track positions (Additional file 2: Fig. S6a). We found that the transition from low to high BrdU content occurred within a very short distance (∼100 bp), indicating that the precision of our estimation of BrdU amount along DNA is in the range of 100 bp. The second source of error comes from replicative track detection. As we run it on BrdU profiles obtained by two methods, TM and CNN, we checked the concordance between them (Additional file 2: Fig. S6b). We found that the median distance between BrdU track starts detected by both methods was 125 bp. We also generated *in silico* chimeric reads for which the transition from low to high BrdU content was known, and we computed the distance between the starting point detected by our algorithms and the theoretical position (Additional file 2: Fig. S6c). We obtained a median distance of 120 and 136 bp using CNN and TM, respectively. Together, these results show that the precision of track start detection by FORK-seq is better than 200 nucleotides.

### Detection of individual initiation and termination events

We then focused on reads containing pairs of divergent or convergent BrdU tracks to map single initiation and termination events (Fig. 3a). We detected 3,606 or 3,877 (2,519 shared) initiation events and 2,953 or 2,784 (1,252 shared) termination events using TM or CNN, respectively. Visual inspection convinced us that events detected with only one of the methods were of similar quality to shared events. We thus pooled all detected (4,964 initiation and 4,485 termination) events for subsequent analyses. Please note that all detected initiation and termination events can be viewed on a web-based interface available at www.biologie.ens.fr/~hennion/forkseq.html. Rapid data examination showed that, as expected, most initiation sites (defined as the midpoints between diverging forks) coincided with known origins [23], ascending RFD segments, and early-replicating peaks on a reported replication timing profile [24], while the termination sites (midpoints between converging forks) were more dispersively located, yet enriched in descending segments of the RFD profile corresponding to late-replicating valleys (Fig. <u>3</u>a, see below for precise quantification). The FORK-seq results were therefore in good agreement with all available cell-population data on the replication program of *S. cerevisiae*. In addition, we observed rare (246/4964) initiation events lacking any known origin (confirmed, likely or dubious origins from OriDB [23]; n = 829) between the diverging forks (Fig. 3b, reads 27, 30, 31, Additional file 2: Fig. S7a,b). Importantly, the visual aspect and the mapping quality scores of these reads were not different from those of initiation events at known origins (Additional file 2: Fig. S7). We consider unlikely that events far from OriDB origins could arise by DNA repair. Nucleotide or base excision or single strand break repair would produce much shorter BrdU tracks (tens of nucleotides at best). Double strand break repair tracks may be longer but should be unidirectional and much rarer than the events here detected. We also observed termination events outside descending RFD segments, of similar quality to events inside these segments (Additional file 2: Fig. S7d; see below).

**Figure 3:**
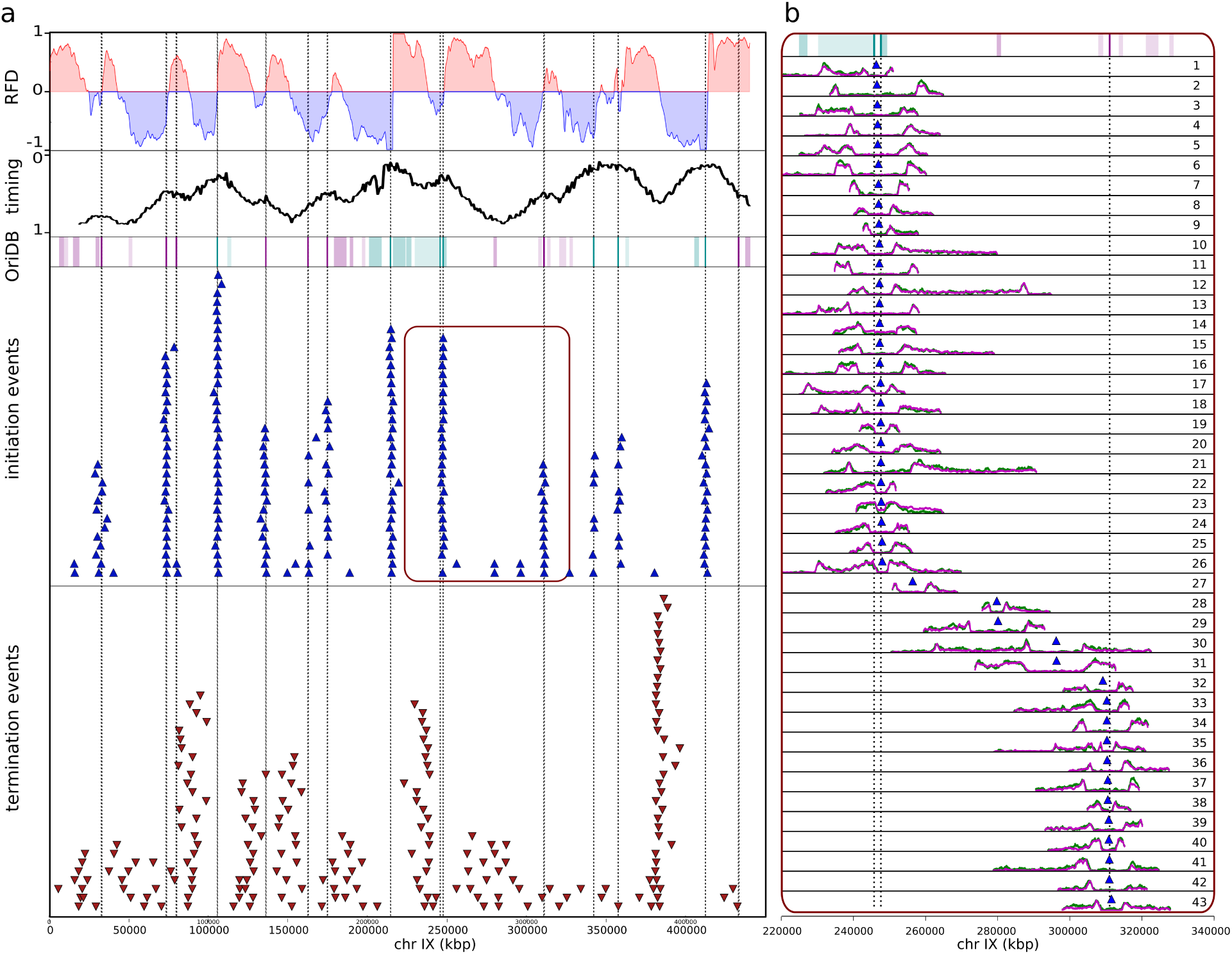
Detection of initiation and termination events. a. Top to bottom: FORK-seq RFD profile obtained merging data obtained from CNN and TM, replication timing profile [24], known origins from OriDB [23] (see Fig.2 legend), detected initiation (blue triangles) and termination (red inverted triangles) events of chromosome IX. Dotted vertical lines: confirmed origins [23]. b. Zoom-in of all initiation events (CNN in pink, TM in green) of the indicated 120 kb segment of chromosome IX (red area).

### Distribution of initiation events in *S. cerevisiae*

A clustering of initiation events is apparent on Fig. 3 (please note that “clustering” is used as per its meaning of “grouping” and should not be confused with the classical “origin clusters” designating multiple initiation events on the same DNA molecule). The proximity of initiation events was objectively assessed by computing the cumulative distribution of all inter-event distances (IEDs) over the genome. The IED distribution was strongly shifted toward short distances compared to a random distribution of the same number of events (Additional file 2: Fig. S8a). We then clustered events at different maximal allowed IED (mIED, Materials and methods) and computed the number of clusters and their dimensions (d, number of events per cluster) as a function of mIED (Fig. 4, Additional file 2: Fig. S8b). As expected, as the mIED increased, the number of isolated events (d = 1) decreased, while the number of clusters (d > 1) first increased then decreased until all events in each chromosome formed a single cluster. This behavior was observed for both the experimental and a control (random) distribution of initiation events. However, the experimental curves differed from the random control by the presence of an extended plateau of stable cluster number at mIEDs of 1-5 kb (Fig. 4a, Additional file 2: Fig. S8b). The distance between the two outmost events in a cluster, or cluster width, also showed a plateau at ∼2 kb over the same mIED range (Fig. 4b), not observed for the random control (Fig. 4d). We concluded that most initiation events tended to cluster within ∼2 kb of each other and that most clusters were separated from each other by > 10 kb. An exemplary chromosomal segment is presented in Additional file 2: Fig. S8b.

**Figure 4:**
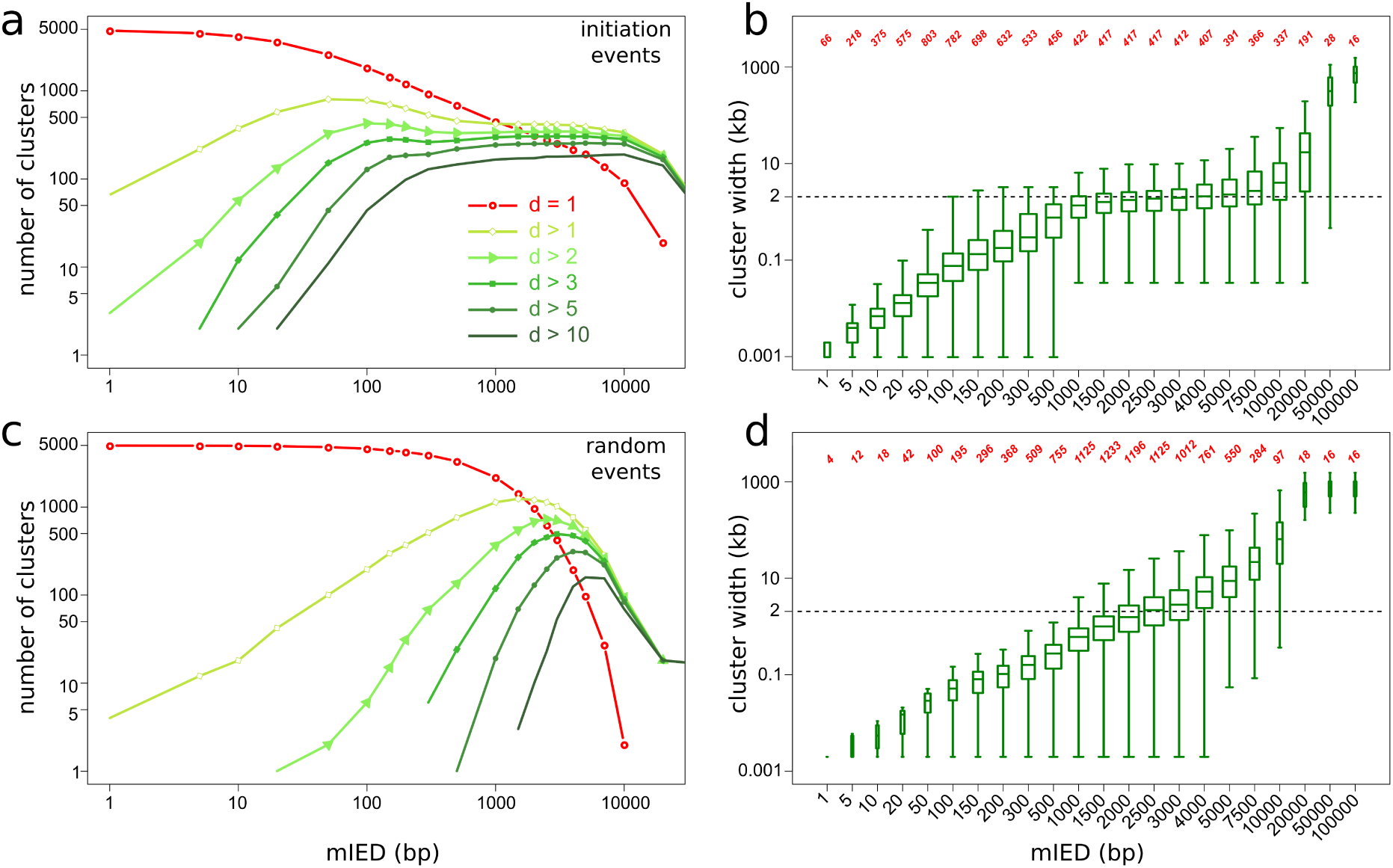
Clustering of individual initiation events. a-d. Number of clusters of indicated dimensions (a,c) and cluster width (d > 1) distribution (b,d) against mIED for experimental (a,b) and randomized (c,d) distribution of initiation events. Box-and-whisker plots (b,d) show median, 2nd and 3rd quartile and range of each distribution. Red numbers (b,d) indicate the number of clusters (d > 1) for each mIED.

Setting the maximal IED at 1.5 kb, 4602 out of 4964 initiation events formed 417 clusters (d > 1) while 362 events remained single (sporadic events; 7.3% of all events, Additional file 1: Table T5). The clusters’ widths ranged from 34 to 7701 bp (median 1579 bp) and their dimension from 2 to 45 (median 7), excluding the rDNA cluster (d=140). 85% of the 417 clusters, but only 35% of sporadic events (extended to 2 kb windows), overlapped with a known origin [23]. Among the OriDB origins, 57% (473/829) overlapped with FORK-seq origins; this fraction was 79% (326/410) for the confirmed OriDB origins, 44% (95/216) for the likely origins and 26% (52/203) for the dubious ones. The overlap was defined after resizing all objects to 2kb. We plotted for clusters of increasing dimension the cumulative distribution of the distance from their median point to the middle of the closest known origin (Fig. 5a). The distance distribution of sporadic events with respect to known origins was not different from random (median distance ∼ 4 kb). In contrast, 69%, 97% and 100% of the clusters of dimension 2-4, 5-12, and > 12, respectively, were ⩽ 2 kb from a known origin center (Additional file 1: Table T5). Therefore, most of the d > 1 clusters coincided with known origins and this coincidence increased with cluster dimension. Only 48 clusters (d > 1) totalling 134 initiation events were located > 2 kb away from known origins (Additional file 1: Table T5, Fig. 3b, reads 30 and 31, Additional file 2: Fig. S7b). These represent weakly preferential initiation sites that escaped detection in previous assays, due to low efficiency (only 4 out of those 48 clusters had d > 4).

**Figure 5:**
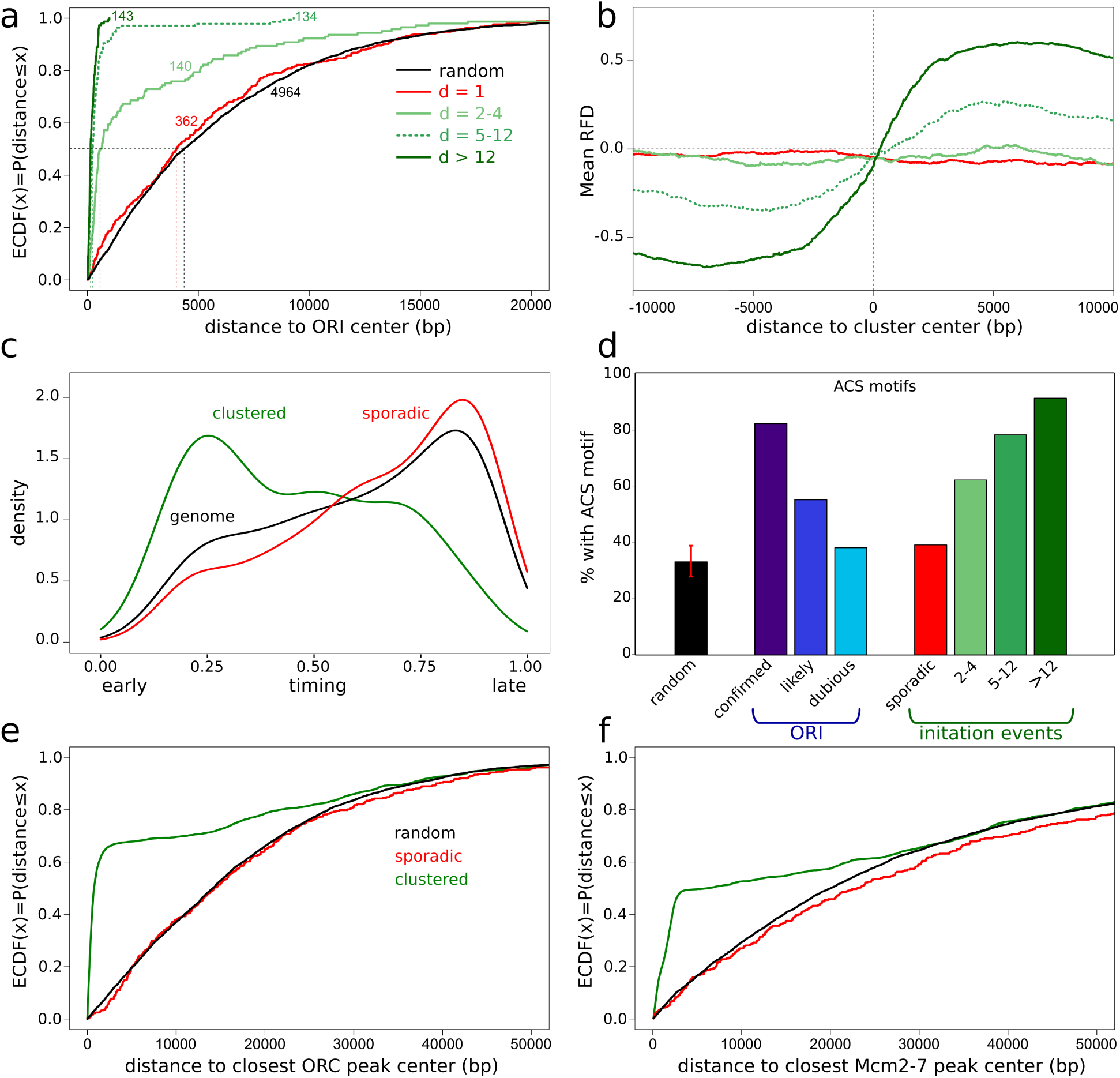
Landscape of individual initiation events. a. Empirical Cumulative Distribution Function (ECDF) of the distances between cluster median points, grouped according to cluster dimension, and the closest known origin (ORI, confirmed, likely and dubious origins from OriDB [23]) center. Black, random elements, red, sporadic initiation events, light, median and dark green, initiation clusters of dimension d = 2-4, d = 5-12 and d >12, respectively. The number of elements in each category is written next to each curve. b. Averaged RFD profile for the clusters grouped according to their dimension (−10kb, +10kb from the cluster median). rDNA cluster was excluded. c. Density distribution of normalized replication timing (0 = start and 1 = end of S phase) for the whole genome (black) and the clustered (green) and sporadic (red) initiation events. d. ACS-motif containing fraction of the indicated elements: randomized initiation events; confirmed, likely and dubious origins from OriDB [23]; sporadic (d = 1) and clustered (d= 2-4; d = 5-12; d >12) initiation events. For randomized elements, the randomization was repeated 1000 times and the median and [0.01,0.99] percentile interval are shown (red bar). e-f. ECDF of the distance between sporadic (red), clustered (green) and randomly shuffled (black) initiation events, and the center of the closest ORC [25] (e) or Mcm2-7 [27] (f) binding site.

Origin efficiency is defined as the fraction of cell cycles in which an origin is active. This parameter can be estimated from cell-population RFD profiles by the amplitude of the RFD upshift at the origin. To check if the number of initiation events per cluster also reflected origin efficiency, we computed mean RFD profiles obtained from CNN data around clusters of increasing dimension (Fig. 5b). As expected, the amplitude of the RFD shifts increased with cluster dimension. The RFD profiles around the sporadic events were null on average, in accordance with their dispersed location in the genome. Origin efficiency has been related to origin firing time. We confirmed that clustered initiation events predominantly occurred in early replicating regions (Fig. 5c). In contrast, the sporadic events followed the same distribution of replication times as the whole genome (Fig. 5c). Altogether, the FORK-seq results show that replication of the *S. cerevisiae* genome combines clustered initiation events at known origins of variable efficiency with a novel class of dispersed initiation events at inefficient sites.

The precision of initiation event detection is similar to the one of individual replicative tracks, the median distance between initiation events detected by TM and CNN methods being 137 bp (Additional file 2: Fig. S6b). To estimate the precision of origin detection, we performed the same clustering procedure independently on the two biological replicates (Additional file 2: Fig. S6d). We obtained 275 clusters common to both replicates and the median distance between their centers was 219 bp. This variability between biological replicates is consistent with the above stated technical precision of track start detection.

### Characterisation of the sporadic initiation events

We assessed whether the sporadic initiation events occurred in a different sequence context than the clustered events. One important feature of the known *S. cerevisiae* origins is the presence of an ARS (autonomously replicating sequence) consensus sequence (ACS). We computed an extended consensus sequence motif from 253 high-confidence origins described in [25] and identified 5,858 genomic sites matching this motif (see Materials and methods). In order to compare objects of similar size, we examined the presence of this motif in a 2 kb window centered on either the midpoint of OriDB origins or the median point of our initiation clusters. The motif was found in 82%, 55% and 38% of confirmed, likely and dubious OriDB origins, respectively; in 91%, 78% and 62% of d > 12, d = 5-12 and d = 2-4 clusters, respectively; in 39% of sporadic events (d = 1); and in 33% of random 2 kb windows (Fig. <u>5</u>d). Thus, the presence of the consensus motif at sporadic initiation events was much rarer than in clusters, and barely more frequent than in random sequences (p-value = 0.03, non-parametric test).

The ACS provides a binding site for ORC. Other required elements specify a nucleosome-free gap that facilitates ORC binding and subsequent loading of the Mcm2-7 complex, a core component of the replicative helicase required for initiation [25, 26]. Using published datasets, we computed the distance between our initiation events and ORC [25] (Fig. 5e) or Mcm2-7 [27] (Fig. 5f) binding sites. As expected, the clustered initation events were found much closer to known ORC and Mcm2-7 binding sites than random genomic positions. On the contrary, the sporadic events behaved very similarly to random sequences (Fig. 5e, f), meaning that there was no detectable enrichment of ORC or Mcm2-7 at these sites. Since no replication can take place in the absence of ORC or Mcm2-7 proteins [28, 29, 30, 31, 32, 33, 34, 35], the low firing efficiency of sporadic origins is likely due to a low efficiency of ORC and Mcm2-7 recruitment rather than to the inhibition of a later origin activation step. This interpretation is reinforced by the lack of sporadic origin enrichment with ACS.

### A two-process model for *S. cerevisiae* genome replication initiation

To summarize, our *ab initio* analysis of 4964 initiation events detected by FORK-seq demonstrated that 93% (4602/4964) were clustered and 7% (362/4964) occurred singly (Additional file 1: Table T5). 95% (4389/4602) of the clustered events, but only 30% (106/362) of the sporadic events, totalling 91% (4495/4964) of all events, were found at ⩽ 2 kb from a known OriDB origin. The remaining 9% (469/4964) of initiation events found > 2 kb away from known origins include 213 clustered and 256 sporadic events, and half (114/213) of these clustered events belonged to low-dimension clusters (d ⩽ 4). All these figures are consistent with a model in which 91% of initiation events coincide with previously known origins, most often clustering in the vicinity of an ACS, while 9% of initiation events occur away from known origins, predominantly dispersively and away from ACSs, forming a novel class of initiation events.

### Distribution of termination events in *S. cerevisiae*

We performed a clustering analysis of termination events detected by FORK-seq as we did for initiation events. Computing the cumulative distribution of all IEDs over the genome, we found that the distribution was shifted toward short distances compared to a random control, but to a lower extent than for initiation events (Additional file 2: Fig. S8a). This is in line with termination events within clusters being more scattered than initiation events as seen on Fig. 3 for the exemplary chromosome IX (see also Additional file 2: Fig. S7d). Setting the maximal IED at 1.5 kb as for initiation events, 3508 out of 4485 termination events formed 785 clusters (d > 1) while 977 events remained single (sporadic events; 22% of all events). The clusters’ widths ranged from 9 to 14,976 bp (median 1,223 bp) and their dimension from 2 to 39 (median 3), excluding the rDNA cluster (d=80). As expected, termination events were located within later replicating regions than initiation events (Additional file 2: Fig. S9a). We computed the mean RFD profiles from combined CNN and TM FORK-seq data around termination clusters of increasing dimension (Fig. <u>6</u>a). As expected, the mean RFD shift was negative around termination clusters, and its amplitude increased with cluster dimension. In contrast, the mean RFD shift was null around sporadic termination events.

The distribution of termination events in *S. cerevisiae* has been previously investigated by cell-population methods [13, 14]. Fachinetti *et al* [13] reported 71 specific termination zones (TERs) defined as regions that are unreplicated late in S phase when replication is slow (*i*.*e*. in the presence of hydroxyurea or at temperatures below 16°C). We found that 66/71 TERs overlapped with FORK-seq termination clusters (Fig. 6b) and that the RFD shift at these TERs was equal to that observed at the strongest clusters (d > 4; Additional file 2: Fig. S9b). McGuffee *et al* [14] used OK-seq to identify 346 descending RFD segments that encompass most (60/71) TERs but are broader. Similarly, we used the previously defined [14] origin efficiency metric (OEM) to measure RFD slope. We identified 341 negative OEM (NOEM), *i*.*e*. descending RFD segments, that encompass 67 TERs but are broader (Fig. 6b). The 341 NOEM segments overlapped 3,630 individual termination events, including 495 within TERs. Importantly, 855 termination events (including 9 within TERs) mapped outside the NOEM segments, *i*.*e*. in positive RFD slopes.

**Figure 6:**
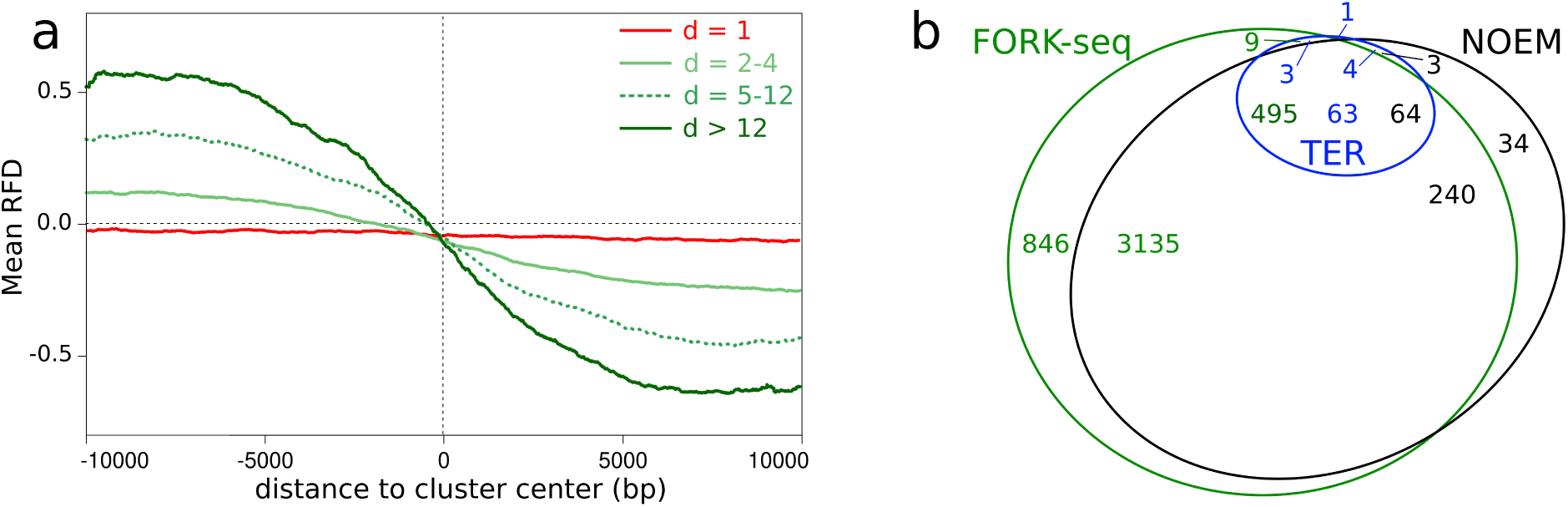
Landscape of individual termination events. a. Averaged RFD profile for the clusters grouped according to their dimension (−10kb, +10kb from the cluster median). rDNA cluster was excluded. Red, sporadic termination events, light, median (dotted line) and dark green, termination clusters of dimension d = 2-4, d = 5-12 and d >12, respectively. b. Venn diagram showing the overlap between FORK-seq termination events, TERs [13] and negative OEM (NOEM) segments. The number of elements in each area are indicated in the same colour as each class of objects.

### Resolving the contributions of initiation and termination events to RFD

The above results suggest that a minority of termination events occur in positive RFD slopes and, reciprocally, that a minority of initiation events occur in negative RFD slopes. To address this point, we used the OEM to measure RFD slopes at individual initiation and termination events (Additional file 2: Fig. S9c, d). We indeed found that 11% of initiation (resp. 18% of termination) events were associated with a negative (resp. positive) OEM. These events were predominantly sporadic: 60% of the sporadic, *vs* 8% of the clustered, initiation events had a negative OEM, and 41% of the sporadic, *vs* 12% of the clustered, termination events had a positive OEM. Therefore, FORK-seq is able to detect a large number of initiation and termination events that are inevitably missed in cell-population analyses. Strictly speaking, the shift in RFD over a considered segment should be proportional to the difference between the number of initiation and termination events over that segment [36]. To verify this point, we computed this difference profile from the 4,964 initiation events and 4,485 termination events. The profile was strikingly similar (Spearmann correlation 0.80) to the OEM profile computed from the assembled RFD profile from the 58,651 oriented BrdU tracks (Fig. 7). Therefore, there is a high consistency betwen the full dispersion of initiation and termination revealed by analysis of individual events and the statistical average of fork direction measured by population analysis of all the oriented BrdU tracks.

**Figure 7:**
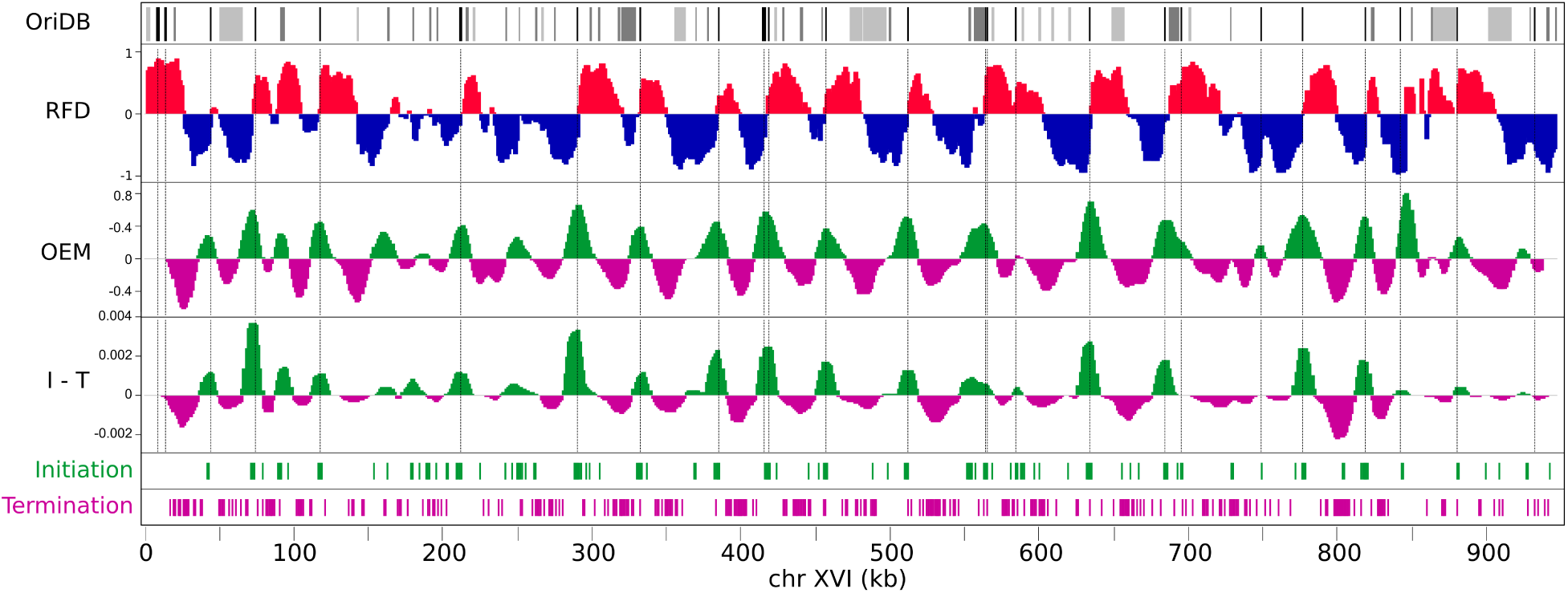
Similarity of initiation minus termination efficiencies computed from the FORK-seq RFD profile or from the individual initiation and termination events. Top to bottom: OriDB origins (black, confirmed, dark gray, likely, light gray, dubious [23]). RFD profile from FORK-seq data; OEM computed from RFD profile; density profile of initiation minus termination events (I-T); individual initiation events; individual termination events.

## DISCUSSION

We have developed a novel method, termed FORK-seq, to map DNA replication genome-wide at the single molecule level, based on *in vivo* replicative incorporation and nanopore sequencing detection of BrdU, a thymidine analog widely used in cell proliferation studies. Several important points made FORK-seq possible. First, nanopore sequencing can read native single DNA strands in the absence of intermediate amplification steps. This allowed direct detection of the *in vivo* incorporated BrdU, revealing the replication pattern and genomic position of each sequenced strand. Second, the use of a thymidine-auxotroph yeast strain with reconstituted thymidine import and salvage pathway allowed efficient BrdU labeling of replication tracks during brief pulse-chase experiments. The short pulse duration (4 min) combined with long read lengths (10-140 kb; Additional file 2: Fig. S10) permitted visualization of mostly complete BrdU tracks along single molecules. Third, we developed and validated against each other two independent methods (CNN and TM) to precisely measure BrdU content variation along the reads, allowing to orient BrdU tracks according to their direction of synthesis. Track orientation was further validated by independent OK-seq data. Fourth, many reads were long enough to contain multiple BrdU tracks, allowing high precision mapping of individual initiation and termination events.

FORK-seq precision was assessed at different levels of analysis. The precision of the transition from low to high BrdU content was ∼ 100 bp. The median CNN-to-TM distance between BrdU track starts and initiation events was 125 bp and 137 bp, respectively. Using chimeric reads, the distance between the detected and theoretical track start was 120 and 136 bp using CNN and TM, respectively. Therefore, the technical precision of FORK-seq is better than 200 nucleotides. The variability of origin detection between two biological replicates, which reflects both the technical imprecision of FORK-seq and the true biological dispersion of initiation events, was 219 bp.

Eukaryotic DNA replication has been best studied in the yeast *S. cerevisiae* [35]. Yeast origins were first identified as ARSs using plasmid replication assays [37]. Physical mapping of replication intermediates in cell populations confirmed that ARSs act as chromosomal replication origins with variable efficiency and failed to detect initiation elsewhere than at ARSs [38, 39, 40, 14, 24]. Genome-wide replication profiles have now attained high spatial (1 kb) and temporal (5 min) resolution and their mathematical analysis suggests that origins fire stochastically and independently of each other [41, 42], as first proposed based on a single-molecule (DNA combing) analysis of yeast chromosome VI replication [43]. A few studies reported that the sequence requirements for yeast origin function can be more flexible than previously thought. Notably, an *in vivo* study showed that replication of an origin-depleted chromosome can initiate from non-canonical sites around deleted origins [44]. *In vitro* reconstitution experiments of plasmid replication using yeast extracts [45, 46] or purified proteins [47, 48] showed that replication can initiate in an ARS-independent manner at high ORC/DNA ratio, although ARS-independent initiation is suppressed by competitor DNA or when nucleosomes are preassembled on the template. Very recently, Müller *et al* developed a nanopore-sequencing based, single-molecule replication mapping method termed D-NAscent, that suggests that up to 20% of all initiation events may occur dispersed through the yeast genome [11]. Using FORK-seq, we report here independent evidence for dispersed initiation, although we find that only 9% of all initiation events occur > 2 kb from a known origin (see below). From the point of view of origin efficiency, only 7% of all initiation events occured singly, and these appear to be randomly dispersed along the genome.

The distribution of termination events in *S. cerevisiae* has been previously investigated by Fachinetti *et al* [13] in the presence of conditions of slowed replication and by McGuffee *et al* [14] by OK-seq in unperturbed cells, but was not further analyzed using D-NAscent. Here, we confirm that termination does occur at the locations previously reported [13, 14] and that TERs are particularly efficient termination regions, but we also find that termination is even more dispersive than previously thought [14], with 18% of termination events located in segments where cell population methods could only detect a predominance of initiation events. This further illustrates the power of FORK-seq for replication profiling.

FORK-seq and D-NAscent are similar in principle but differ in details. Detection of BrdU in D-NAscent relies on the identification of thymidine-containing hexamers where the distribution of electrical currents is significantly affected by the presence of BrdU, allowing assessment of BrdU incorporation on average every 21 nucleotides in the yeast genome. This is related to RepNano TM strategy where we identified pentamer-to-pentamer transitions whose current shift was significantly affected by BrdU, allowing assessment of BrdU content every 18 nucleotides on average. RepNano second, CNN-based, strategy does not attempt to call individual bases but estimates BrdU content for 96-nucleotide sliding windows. Importantly, CNN estimates nicely correlate with TM-based measurements. The BrdU signal obtained by D-NAscent software detection was located at similar positions to RepNano but was weaker, precluding automatic BrdU track detection and orientation and assembly of a valid RFD profile. The number of B calls was about 4 times lower in D-NAscent than in TM: in the highly substituted sample, TM and D-Nascent called 10.3% and 2.6% of all T sites, respectively. This represents a significant difference between the D-NAscent and FORK-seq computational methodologies. The other significant difference between the DNAscent and FORK-seq experiments is the replication labeling strategy. Müller *et al* labelled replication by synchronizing thymidine-prototroph cells in G1 and releasing them in the presence of limiting BrdU concentrations. They found that the level of BrdU incorporation decreased as S phase progressed, suggesting that the intracellular ratio of BrdUTP to dTTP declined due to increasing activation of endogenous thymidine synthesis and/or declining rate of BrdU import. They measured gradients of BrdU incorporation along single reads by computing a z-score of BrdU incorporation by non-overlapping 2-kb windows and used these gradients to deduce replication direction and to map initiation events. In contrast, our FORK-seq experiments were carried out using asynchronously growing, thymidine-auxotroph cells subjected to a brief BrdU pulse followed by a chase with a 10-fold excess of thymidine. This strategy allowed a much more precise control of the intracellular ratio of BrdUTP to dTTP and allowed us to map the start of BrdU tracks with a ∼200 nucleotide resolution, a much better precision than D-NAscent. In addition, we found a more even distribution of early and late initiation events than Müller *et al* (Additional file 2: Fig. S11a). Our interpretation is that in D-NAscent experiments, the change in the intracellular ratio of BrdU to thymidine is steeper in early than in late S phase due to the use of synchronized cells, whereas in FORK-seq experiments there is an unbiased representation of replication tracks regardless of replication time due to the avoidance of cell synchronization.

We found a tighter distribution of initiation events around known origins (Additional file 2: Fig. S11b; 91% of all events at ⩽ 2 kb from a known origin, median distance 434 bp) than Müller *et al* (57% at ⩽ 2 kb, median distance 1,746 bp). This result likely comes from the better spatial precision of FORK-seq which can explain in part the discrepancy between the two studies regarding the fraction of initiation events judged to map significantly away from known origins (20% at > 3.9 kb vs. 9% at > 2 kb). It is also possible that the different genetic background of the thymidine-prototroph and auxotroph strains, and the different synchronization and labelling procedures, affect the prevalence of dispersive *vs* site-specific initiation by perturbation of dNTP pools and fork progression or by other mechanisms. Further studies are required to resolve these questions. This discrepancy left aside, both studies demonstrate the power of nanopore sequencing for replication studies and reveal the existence of dispersed replication initiation throughout the yeast genome. In addition, we provide evidence that dispersed initiation events are barely if at all enriched in ACS motifs typical of site-specific initiation events and inefficiently recruit ORC and Mcm2-7 complexes, and that termination is more dispersive than previously thought. Together with previous *in vitro* [45, 46, 47, 48] and *in vivo* [44] studies, these results indicate that many genomic locations can function as replication origins at low frequency. The superposition of efficient initiation at few specific sites with inefficient initiation at widely dispersed sites is reminiscent of the strategy proposed for mammalian genome replication [22] and leads to a more unified view of eukaryotic DNA replication.

FORK-seq relies on the incorporation of BrdU in replicating DNA, which is achievable in many experimental systems. All the sequencing reads used in this study were obtained with only six runs on the MinION sequencing apparatus, and we now routinely obtain the same number of reads with one sequencing run. When applied to mammalian cells, with larger genomes and wider spaced origins, both the length and number of sequenced DNA molecules may become limiting. However, read lengths of up to 2.3 Mb have been reported [6] so there is no absolute technological limitation to improve read length. As to throughput, the recent availability of the PromethION, which delivers 10 times more data per single flow cell and can run 48 flow cells in parallel, should allow high coverage studies of human genome replication in the near future.

## CONCLUSIONS

This work demonstrates the power of nanopore sequencing to study DNA replication at the single-molecule level following replicative incorporation of BrdU. FORK-seq allows to reproduce genome-wide RFD profiles and to rediscover *ab initio* the known replication origins and termini of *S. cerevisiae*. In addition, it maps low efficiency origins as well as dispersive initiation and termination events undetectable by population methods. This high resolution genome-wide method represents a major step forward compared to current single-molecule methods and will likely transform this research field.

## MATERIALS AND METHODS

### Synthetic template construction

pTYB21 plasmid (New England Biolabs, NEB) was linearized with EcoRV (NEB) and purified using home-made SPRI (Solid Phase Reversible Immobilization) beads. After initial denaturation for 5 min at 94°C, primer extension was performed on 200 ng of linear plasmid, using 0.5 U/μL LongAmp Taq DNA polymerase (NEB) with 300 μM of each dNTP (with either dTTP or BrdUTP (Thermo

Fisher Scientific, TFS)) and 400 nM of NanoP-pTYB-F primer (5’-ATCGTCGACGGATCCGAATTCCCTGCAGGTAATTA CGGCTGCTAACAAAGCCCGAAAGGAAGCTGAGTTGGCTGCTGCCACCGCT-3’; Eurofins) for 20 min at 65°C. A control (‘water’ lane in Fig. 1b) with only dATP, dGTP and dCTP was also included. The ssDNA was then digested by ExoSAP iT (TFS) for 30 min at 37°C and the DNA was purified using SPRI beads. The size of the product was assessed using an Agilent DNA 12000 chip on a Bioanalyzer and its amount was quantified using Qubit dsDNA HS Assay (TFS).

### Samples used for CNN learning

MCM869 genotype is *MATa ade2-1 trp1-1 can1-100 leu2-3 his3-11,15 ura3-1::URA3-GPD-TK7x aur1::AUR1-C-ADH-hENT1 bar1::LEU2 cdc21::kanMX* [20]. PSL genotype is *MATa trp1-1 leu2-3,112 his3-11,15 ura3-1::URA3-GPD-TK aur1::ADH-hENT1 bar1::LEU2*. MCM869 and PSL were grown at 30°C in YPD with 100 μM thymidine (Sigma-Aldrich). We used 7 samples for the training of the CNN (see Additional file 1: Table T3): two negative controls with MCM869 grown in 100 μM thymidine (Thy_R_1_ and Thy_R_2_), two samples with intermediate BrdU amount prepared from PSL strain (CDC_21_ proficient) grown for 24h in 100 μM (BrdU_9) or 1 mM (BrdU_38) BrdU, one sample with intermediate BrdU amount obtained from MCM869 grown in a mixture (75:25) of BrdU and thymidine for 24h (BrdU_69), and two highly substituted samples from MCM869 grown in 100 μM BrdU for 24h (BrdU_R_1_ and BrdU_R_2_). The naming follows the BrdU content as measured by mass spectrometry (Additional file 1: Table T_3_; see below).

### Pulse-chase labelling

MCM869 grown for 3h in fresh YPD medium with 100 μM thymidine, were washed and transferred in YPD for 30 min, pulsed with 100 μM BrdU for 2 min, added with 33 μM thymidine for 2 min, chased with 1mM thymidine for 45 min, pelleted and frozen. DNA was purified by Zymolyase, RNAse A and proteinase K digestion followed by Qiagen Genomic-tips according to the manufacturer instructions. The size of the DNA was checked by agarose gel electrophoresis and on an Agilent TapeStation.

### MCM869 OK-seq

Freshly inoculated MCM869 cells in 10 L of YPD + 100 μM thymidine were grown to an OD600 of 0.7 (200 × 10^9^ cells), pelleted at 4,000 g at 30°C for 10 min, resuspended and shaken in 8 × 100 mL flasks of prewarmed YPD + 100 μM 5-ethynyl-2’-deoxyuridine (EdU) for 2 min, added with 2.5 mL of 0.5 M EDTA per flask, chilled on an ice-water bath and pelleted at 4,000 g for 10 min at 4°C. Cells were rinsed twice with ice-cold water, resuspended at 1.5 − 2 × 10^9^/mL in ice-cold nuclear isolation buffer (NIB; 17% glycerol, 50 mM MOPS, 150 mM potassium acetate, 2 mM magnesium chloride, 500 μM spermidine, and 150 μM spermine, pH 7.2), dispensed in 50 mL plastic centrifuge tubes (<13 mL of cell suspension per tube), mixed with an equal volume of glass beads (0.45-0.52 mm diameter), subjected to 15 cycles of high speed vortexing for 30 s followed by > 30 s incubation on ice in a 4°C cold room. The supernatant was collected and mixed with two rinses of the beads with NIB and pelleted at 5,500 g for 10 min at 4°C to collect nuclei. Subsequent DNA extraction, heat denaturation, size-fractionation on sucrose gradients and Okazaki fragment purification and sequencing were as described previously for human cells [22] with the following minor modifications. Biotinylation was conducted with 0.1x PBS pH 7.2, 2 mM CuSO4 freshly pre-mixed with 10 mM Tris((1-hydroxy-propyl-1H-1,2,3-triazol-4-yl)methyl)amine (THPTA) and 10 mM sodium ascorbate for 45 min in dark. To maintain a high ratio of DNA to streptavidin beads, only 100 μg Dynabeads MyOne Streptavidin T1 were used to capture the biotinylated Okazaki fragments.

### Mass spectrometry

LC-MS/MS was performed using a TSQ Quantiva triple quadrupole mass spectrometer (Thermo Scientific) coupled to an UltiMate 3000 XRS HPLC system (Dionex, Thermo Scientific). Metabolites were quantified with mass spectrometry, using selected reaction monitoring (SRM) in the positive ion mode. In source generated ions were used as quantifiers using the following SRMs (dT m/z 127 to 54; BrdU m/z 191 to 118 and m/z 193 to 120). 1 μg of genomic DNA was digested by 10U of DNA Degradase Plus (Zymo Research) overnight at 37°C. Before analysis, formic acid in water was added to 0.1%. Metabolites were separated using an 8 minutes gradient starting at 1% mobile phase A (0.1% formic acid in water) and ramping up to 60% A in phase B (0.1% formic acid in acetonitrile) using a C18 reversed phase column (Kinetex 2.6 μm C18 100 Å, LC Column 150 × 2.1 mm; Phenomenex) and employing a flow rate of 100 μL/min. Samples were analyzed in technical duplicates and MS data were analyzed using the software TraceFinder (Thermo Scientific). A defined mixture of BrdU and dT was used for measuring the combined relative ionization and fragmentation efficiencies of these ions used for calculating the percentage of BrdU incorporation at native thymidine sites.

### Nanopore sequencing and data preprocessing

MinION sequencing libraries were prepared according to the manufacturer protocols using the 2D low input kit with R9 chemistry for the plasmid experiments and the 1D ligation kit with R9.5 or R9.4.1 chemistry for the yeast experiments. ONT protocol and software versions used for the different samples are listed in Additional file 1: Table T1, T3 and T4.

For the plasmid experiments, raw reads were basecalled using Metrichor (ONT) with default parameters and aligned on the reference using BWA MEM [19] with the -x ont2d option (−k14 -W20 -r10 -A1 -B1 -O1 -E1 -L0). The event sequences were extracted using poretools [49] and aligned on the reference using the sam file information and custom Python scripts. Current values were normalized by subtracting the whole read average current. The PCA plot was generated with R *pcaMethods* library using as inputs 1kb long current sequences (positions 100-1100 on the reference plasmid sequence) from 1000 reads of each sample.

For the yeast experiments, raw reads were basecalled using Albacore (ONT). We used Tombo (ONT) *preprocess annotate_raw_with_fastqs* to add the DNA sequence obtained from Albacore into the raw fast5 file. The sequence was then mapped and corrected using the reference genome and the current signal was segmented and assigned to each base using Tombo *resquiggle* command (which calls minimap2). These two steps output a sequence of bases associated with its average current (N_i_, m_i_). Next steps are integrated in the RepNano software (available at https://github.com/organic-chemistry/RepNano [21]). RepNano first computed the current shifts δ_i_ = m_i_ − m_i+1_ and used (N_i_, δ_i_) as the input of further processing steps.

### Current shift normalization and computation of reference transition matrices

In order to remove possible drifts in current values between reads, we defined a two-step normalization and filtering procedure as follows. Using 4000 reads of a given experiment, we computed the average current shift value associated to each possible pentamer-to-pentamer transition (i.e. for each hexamer N^(6)^) and obtained a first transition matrix, TM, such as 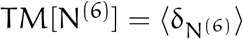 (step1). Second, for each read, this matrix was used to determine the sequence of expected current shifts 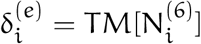. We determined the drift for each read by determining the linear transformation of the experimental signal (αδ_i_ + β) that minimized the difference with the expected signal disregarding transition with a T in the center of either pentamer 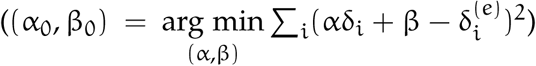. Reads where the quadratic optimized distance was greater than 0.25 were discarded. Finally, the dataset was updated with the drift normalized current shift signals obtained for each read: 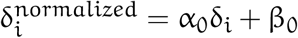 (step 2).

We applied this procedure twice on the two thymidine samples (Thy_R1 and Thy_R2), and the two BrdU rich samples (BrdU_R1 and BrdU_R2), removing 4% of the thymidine sample reads and 12% of the BrdU sample reads. We then averaged the 2 resulting thymidine (resp. BrdU) transition matrices to determine the matrix TM_T_ (resp. TM_B_) containing the expected normalized current shift for transition in the thymidine and BrdU context 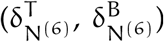, respectively. In the same manner, we applied the second step of the normalization procedure using TM_T_ as the reference transition matrix to all samples resulting in homogeneously normalized data used in all subsequent analysis steps. At this point, each read was represented by a sequence 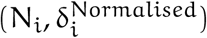.

### BrdU content determination using transition matrices

As expected from the *in vitro* experiment (Additional file 2: Fig. S1d), the difference between TM_T_ and TM_B_ for transition involving a pentamer having a T in its middle position was much larger than for the other transitions (Additional file 2: Fig. S3d); 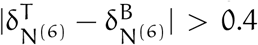 being (514 transitions) almost exclusive of transitions involving pentamers with a T in the middle position (507 transitions). Hence, we used the following strategy to call B or T at T sites (considering the hexamers where Ts are in position 3). We looked up in TM_T_ and TM_B_ for the expected current shift for the site: 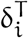 and 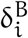. If 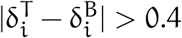, we called a T or a B according to the shortest distance between the observed δ_i_ and either 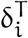 or 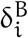. If the difference was ⩽ 0.4, we called the indeterminate base a X.

### Neural network architecture

RepNano CNN architecture was designed so as to directly estimate BrdU content in a given window thus avoiding calling individual incorporation of BrdU. This approach entails dimensionality reduction from the window size to 1. This is achieved by the action of several layers each performing a fraction of the required dimensionality reduction. We therefore implemented a 1D convolutional neural network (Additional file 2: Fig. S4) to predict the average BrdU content for a segment of 96 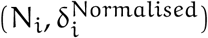 plateaus (see below). Each base was encoded using one hot encoding (A = (1,0,0,0), T = (0,1,0,0), C=(0,0,1,0), G=(0,0,0,1)). Each plateau was converted to a 5-dimensional vector, where the four first dimensions accounted for the base in the one-hot encoding scheme and the last one was 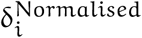. This gave an input of shape (96,5). The neural network consisted in three convolutional layers with 32 filters of kernel size 5. After each of the two first convolutional layers, we used a layer of maxpooling over 4 consecutive values that each reduces the size of the vectors by a factor of 4, so that their combined action is a 16-fold dimensionality reduction. This implies the input window size to be a multiple of 16. Moreover it must contain enough informative transitions to reliably train the neural network. We evaluated using the TM method that 514 (12.5%) of the 4,096 pentamer-to-pentamer transitions were informative for BrdU incorporation. The CNN window size was therefore chosen to be 96 bp as this is a multiple of 16, containing 14 informative transitions on average, which we reasoned was enough. After the third convolutional layer we obtained a vector of shape (6,32). On the first dimension we applied a dense network with a sigmoid activation and one output leading to a vector of shape (6,1), that was then averaged on the first dimension. CNN was implemented in Python with the Keras library [50].

### CNN learning

We used 4000 reads of each of the 7 samples described above for CNN learning. The learning was done splitting the mapped reads into sequences of length 96, and the error between the prediction and the correct sequence was minimised using a stochastic gradient descent optimizer implemented in Keras [50] using default parameters, a learning rate of 0.1, a decay rate of 10^−6^ and a momentum of 0.9. The learning dataset was randomly split in 90% training and 10% validation sequences. After each epoch the loss was evaluated on the validation set and after 5 consecutive steps with no improvement of the validation loss, the learning was stopped. We then computed the average BrdU content for the reads coming from the BrdU-rich samples, which allowed us to separate BrdU-substituted reads from parental thymidine reads. We then used this cleaned dataset to perform the learning again.

### CNN calling

RepNano uses the CNN to predict for each read the BrdU content by 96-nucleotide overlapping windows shifted by 10 bases. Then the average BrdU content is computed per 10-nucleotide segment by averaging on the 96-nucleotide windows containing this segment.

### D-NAscent calling

We downloaded D-Nascent from https://github.com/MBoemo/DNAscent (on 2019/11/20). After compilation, we used the *detect* function to call BrdU from our fast5 files. As described in [11], we called a BrdU when the log-likelihood was > 2.5 and we used as input for subsequent track detection the number of B per 100 bp windows.

### BrdU track calling and orientation

A home-made Python script was used to detect and orient replication tracks. It is included in the RepNano package. Reads < 5 kb were not used for this analysis. For CNN and TM, BrdU content along the reads was smoothed over 30 consecutive positions corresponding to a B or a T after TM basecalling (present on average every 18 bases). For D-NAscent, we smoothed on two 100 bp-windows. We discarded reads with no tracks by filtering on both a minimal standard deviation of the BrdU proportion on the whole read (0.115 for CNN, 0.3 for TM, 0.02 for D-Nascent), and a minimal signal amplitude (0.4 for CNN, 0.5 for TM, 0.08 for D-Nascent). The signal was rescaled between 0 and 1 and subjected to piecewise linear segmentation using the Ramer–Douglas–Peucker algorithm implemented in the Python module *simplification*. A track was detected when an upward shift (amplitude > 0.44) was followed by a downward shift (amplitude > 0.26). The track was then oriented according to the different shift slopes (−slope_1_/slope_2_ > 1.5) or BrdU content of the preceding and following minima (min_f_ − min_p_ > 0.075; the level of BrdU decays during the chase but remains higher than before the pulse). A bed file containing the coordinates of all oriented tracks was output and used to compute RFD profiles. We added two scores quantifying the fork asymmetry (Ascore) and the amplitude of the starting jump (Jscore) for subsequent filtering. Initiation (termination) events were determined by the presence of two consecutive high confidence (|Ascore| > 2 and Jscore > 1) diverging (converging) tracks. In addition, a minimal gap of 1 kb between fork starts and a maximal value (0.14 for CNN, 0.12 for TM) for the plateau (mean value) between the forks were required for initiation calling. We imposed the same maximal value for the fork starts to call termination events. Events were considered as shared between CNN and TM if the midpoints between fork starts obtained by the two methods were within 2 kb.

### RFD profiles

All following procedures are presented in the script RepNanoScript.r. CNN and TM fork data were imported into R as *GenomicRanges* aligned to the SacCer3 version of the yeast genome. OK-seq data from our lab were imported in R as *GenomicAlignments* and paired-end reads were converted into *GenomicRanges*. Duplicates were removed at this step. OK-seq data from the ligase mutant [14] were downloaded from NCBI GEO website (GSE33786) and also converted into *GenomicRanges*. RFD was computed using R (see RepNanoFunction.r script) as the difference between rightward- and leftward-fork coverage normalised by total coverage. The data were smoothed using a running mean with a 1 kb window. RFD profiles were exported into bigwig files using the export function from the *rtracklayer* package. Spearman correlations between RFD profiles were computed with the base R *cor* function and a heatmap was produced with *ggcorplot* package.

### Analysis of initiation and termination events

Initiation events defined as the midpoints between diverging forks obtained from CNN or TM were merged and imported into R as *GenomicRanges*. Known origins were imported from OriDB 2.1.0 [23](http://cerevisiae.oridb.org/) into *GenomicRanges* objects. Inter-event distance and distance to the nearest origin center were computed using the *distanceToNearest* function from the *GenomicRanges* package. Empirical Cumulative Distribution Function (ECDF) was computed using the *Ecdf* function from *Hmisc*. For control analyses, initiation events were randomly redistributed by shuffling them while keeping constant the number of initiation events per chromosome. The randomized events were subjected to the same analyses as the experimental events. Clustering of initiation events was performed by sorting events by chromosomal order and computing the next neighbour distance. Neighbouring events closer than the clustering limit (mIED) were assigned to the same cluster. The width of a cluster was defined as the distance between its outmost elements, and its dimension d as its number of events. For further analyses, the mIED was set to 1.5 kb. Clusters were binned according to their dimension into isolated events (d = 1) and three groups of similar size (d = 2-4, d = 5-12 and d > 12, ∼ 130 clusters in each group). ECDF was computed with the distances from the cluster median point to the nearest known origin center for each cluster. ECDF for each group was then compared to the global ECDF of the cluster of shuffled events. Mean RFD profiles from Fig. 5b and 6a were computed using RFD calculated with oriented forks from FORK-seq merged (CNN and TM) data and centered on the median position of clusters. Replication timing data was taken from [24] using the *GSM1180749_T9475_Illumina_normalised*.*wig* file, lifted over from SacCer1 to SacCer3 version of the genome, binned in non-overlapping 1kb windows and normalised between 0 and 1 (start and end of S phase, respectively). The timing density plot shown on Fig. 5c was created using the *densityplot* function from the *lattice* R-package. The position weighted matrix (PWM) for the ACS was computed by realigning the ORC-ACS coordinates from [25] with *AlignSeqs* function from *DECIPHER* R-package. The PWM matrix was generated with the *seqLogo* R-package. The genomic map of all positions matching this PWM (score > 80%) was generated with *matchPWM* from *BioStrings* package and compared to known origins and clusters positions. To perform the random control, coordinates from the 362 isolated events were shuffled 1000 times, and the median and [0.01,0.99] percentile interval were computed. ORC and Mcm2-7 position were extracted from published data and imported as *GenomicRanges* in R (*GSM424494_wt_G2_orc_chip_combined*.*bed* [25] and *GSM932790_Hu-MCM-Comb-Norm-Peaks-WT*.*bed* [27], respectively). Distance to their centers and ECDF were computed as previously described. TER data were manually copied from the supplementay table S3 in [13] and imported in R as *GenomicRanges*. Euler plot for Fig.6b was generated using *EulerAPE_3*.*0*.*0*.*jar* application (http://www.eulerdiagrams.org/eulerAPE/) using intersection provided by the *overlapsAny* function from the *GenomicRanges* package. OEM (Origin Efficiency Matrix) were computed as described in [14] from the FORK-seq oriented forks on 10 kb overlapping windows. NOEM corresponds to area of strictly negative OEM. D-NAscent called origins were imported from GEO data (*GSM3450332_1x_BrdU_final*.*calledOrigins*.*bed*). Distance to OriDB origins centers was computed as described before using the middle of D-NAscent called origins. Replication timing for D-NAscent origins, FORK-seq initiation and termination events was affected as described before (replication timing data from [24]). The density profile of initiation minus termination events shown on Fig.7 was computed by binning initiation minus termination events in non-overlapping 5 kb windows and smoothed using a 10 kb running mean.

## DECLARATIONS

### Availability of data and material

The datasets generated during the current study are available from the ENA repository [PRJEB36782] [51]. Raw data, as well as bed files with replication forks, initiation and termination events are included. The reads containing the 4,964 initiation and 4,485 termination events can be visualized at www.biologie.ens.fr/~hennion/forkseq.html. RepNano can be downloaded from https://github.com/organic-chemistry/RepNano or https://doi.org/10.5281/zenodo.3743241 [21].

## Author’s contributions

OH designed the project and raised fundings. MH, FP, BT, XW and BLT performed the experiments. CC, SE and AL performed the nanopore sequencing. JMA and BA implemented RepNano. MH, LL and EN analysed the data. MH and OH wrote the manuscript with inputs from the other authors.

## Acknowledgements

This article is dedicated to the memory of Alain Arneodo, who passed away during its writing. The authors acknowledge Alain’s enthusiam and constant support. The authors also thank all members of the OH lab as well as Arach Goldar, Patrice Abry and Nelly Pustelnik for helpful discussions. We thank Erwan Sallard for his help generating the PSL strain. JMA and BA thanks BioSyL Federation and Ecofect LabEx [ANR-11-LABX-0048] for inspiring scientific events. This work (OK-seq experiments) has benefited from facilities and expertise of the High-throughput Sequencing Platform of I2BC (Gif-sur-Yvette). We thank especially Yan Jaszczyszyn for expert help with Okazaki fragment sequencing. We thank the sequencing facility of François Jacob Institute of biology (Genoscope, Evry) for performing MinION sequencing. Finally we thank Michael Mitter and Vienna BioCenter Metabolomics facility for mass spectrometry measurements.

