## Additional file 1: Tables for "FORK-seq: replication landscape of the *Saccharomyces cerevisiae* genome by nanopore sequencing"

| Sample ID | dTTP | BrdUTP |
|---|---|---|
| Run ID | 160809_MN18874_FAD19767 | 160809_MN17490_FAD19641 |
| Library protocol | 2D-MAP006 low input | 2D-MAP006 low input |
| DNA used for library (ng) | 120 | 100 |
| Flowcell | R9 | R9 |
| MinKNOW version | 1.0.2.0 | 1.0.2.0 |
| Basecaller version | 2.40.17 | 2.40.17 |
| Workflow | 2D RNN for NSK007 | 2D RNN for NSK007 |
| Metrichor run ID | 113694 | 113693 |
| Number of reads | 114750 | 77100 |
| Number of "pass" reads | 20583 | 10328 |
| Number of "fail" reads | 94167 | 66772 |
| % mapped | 60.0 | 49.2 |

Table T1: Detailed information about *in vitro* template sequencing runs. The "% mapped" is the percentage of reads mapped by BWA MEM after Metrichor basecalling.

| | dTTP | | | BrdUTP | | |
|---|---|---|---|---|---|---|
| | pass | fail | all | pass | fail | all |
| Number of reads | 20583 | 94167 | 114750 | 10328 | 66772 | 77100 |
| % of pass reads | | | 17.9 | | | 13.4 |
| TP/CP pairs (2D reads) | 20583 | 47030 | 67613 | 10328 | 33471 | 43799 |
| % of 2D reads | 100.0 | 49.9 | 58.9 | 100.0 | 50.1 | 56.8 |
| TP and/or CP unmapped | 293 | 32338 | 32631 | 113 | 25414 | 25527 |
| TP/CP pairs, at least one mapped | 20317 | 23814 | 44131 | 10254 | 13634 | 23888 |
| TP/CP pairs, both mapped | 20290 | 14692 | 34982 | 10215 | 8057 | 18272 |
| TP mapped on F | 10305 | 23357 | 33662 | 4305 | 11932 | 16237 |
| TP mapped on R | 10009 | 29544 | 39553 | 5946 | 17850 | 23796 |
| CP mapped on F | 9987 | 7879 | 17866 | 5914 | 4636 | 10550 |
| CP mapped on R | 10306 | 8013 | 18319 | 4304 | 4604 | 8908 |
| TP unmapped | 269 | 41266 | 41535 | 77 | 36990 | 37067 |
| CP unmapped | 290 | 31138 | 31428 | 110 | 24231 | 24341 |
| TP + CP | 41166 | 141197 | 182363 | 20656 | 100243 | 120899 |
| TP + CP mapped | 40607 | 68793 | 109400 | 20469 | 39022 | 59491 |
| TP + CP unmapped | 559 | 72404 | 72963 | 187 | 61221 | 61408 |
| % mapped | 98.6 | 48.7 | 60.0 | 99.1 | 38.9 | 49.2 |

Table T2: Mapping summary for the *in vitro* templates. TP, template strand (first strand to enter the pore), CP, complement strand (second strand). The mapping was performed independently for the two strands, using BWA MEM. F, forward plasmid strand (primer extended), R, reverse strand (parental).

| Sample ID | Thy_R1 | BrdU_R1 | Thy_R2 | BrdU_R2 | BrdU_9 | BrdU_38 | BrdU_69 |
|---|---|---|---|---|---|---|---|
| Run ID | MN19358_FAH14273 | MN17490_FAH14319 | FAH58492_MN1747 | FAH54284_MN19358 | MN17479_FAK06634 | MN17273_FAK06596 | FAH58543_MN19358 |
| Sequencing date | 20170808 | 20170821 | 20180214 | 20180213 | 20181010 | 20181010 | 20180220 |
| DNA used for library (ng) | 1507 | 1502 | 2022 | 2200 | 1499 | 1505 | 1941 |
| Flowcell | R9.5 | R9.5 | R9.4.1 | R9.4.1 | R9.4.1 | R9.4.1 | R9.4.1 |
| MinKNOW version | 1.7.10 | 1.7.14 | 1.10.23 | 1.10.23 | '1.14.1 | '1.14.1 | 1.10.23 |
| Albacore version | 1.2.3 | 1.2.3 | 2.1.10 | 2.1.10 | 2.3.3 | 2.3.3 | 2.1.10 |
| Number of reads | 92928 | 502122 | 93697 | 606184 | 161954 | 145595 | 367216 |
| % Tombo + RepNano | 62.1 | 54.3 | 87.9 | 59.7 | 78.0 | 70.4 | 83.4 |
| BrdU % (mass spec) | - | - | 0 | 91 | 9 | 38 | 69 |

Table T3: Detailed information about yeast training sample sequencing runs. Sequencing date is in YYYYMMDD format. The "% Tombo + RepNano" is the percentage of reads successfully processed by Tombo and RepNano.

| Sample ID | pulse_R1_1 | pulse_R1_2 | pulse_R1_3 | pulse_R2_1 | pulse_R2_2 | pulse_R2_3 |
|---|---|---|---|---|---|---|
| Run ID | FAH59201_MN17490 | MN17490_FAH68914 | MN18874_FAH68992 | FAH58519_MN18874 | MN19040_FAH69011 | MN19358_FAH66366 |
| Sequencing date | 20180226 | 20180416 | 20180416 | 20180226 | 20180416 | 20180416 |
| DNA used for library (ng) | 2000 | 1518 | 1518 | 1999 | 1736 | 1736 |
| Flowcell | R9.4.1 | R9.4.1 | R9.4.1 | R9.4.1 | R9.4.1 | R9.4.1 |
| MinKNOW version | 1.10.23 | 1.11.5 | 1.11.5 | 1.10.23 | 1.11.5 | 1.11.5 |
| Albacore version | 2.1.10 | 2.1.10 | 2.1.10 | 2.1.10 | 2.1.10 | 2.1.10 |
| Number of reads | 275030 | 255596 | 388174 | 354300 | 224498 | 191502 |
| % Tombo + RepNano | 78.7 | 84.2 | 86.3 | 83.8 | 86.1 | 84.9 |

Table T4: Detailed information about pulse-chase sample sequencing runs.

| | number of initiation events | | | | | number of clusters | | |
|---|---|---|---|---|---|---|---|---|
| | all | events ORI $\leq$ 2kb | events ORI >2kb | cluster median ORI $\leq$ 2kb | cluster median ORI >2kb | all | cluster median ORI $\leq$ 2kb | cluster median ORI >2kb |
| $d$=1 | 362 | 106 | 256 | 106 | 256 | 362 | 106 | 256 |
| $d$=2-4 | 373 | 259 | 114 | 263 | 110 | 140 | 96 | 44 |
| $d$=5-12 | 1060 | 1012 | 48 | 1036 | 24 | 134 | 130 | 4 |
| $d$>13 | 3169 | 3118 | 51 | 3169 | 0 | 143 | 143 | 0 |
| $d$>1 | 4602 | 4389 | 213 | 4468 | 134 | 417 | 369 | 48 |
| total | 4964 | 4495 | 469 | 4574 | 390 | 779 | 475 | 304 |

Table T5: Summary of the clustering of initiation events. Left, initiation events; right, clusters. For each indicated cluster dimension $d$, the given numbers from left to right are those of: all events; events close to ($\leq$ 2 kb) or far from ($>$ 2 kb) a known origin (ORI); events whose cluster median point is close to ($\leq$ 2 kb) or far from ($>$ 2 kb) a known origin (ORI); all clusters of indicated dimension; clusters of indicated dimension whose median point is close to ($\leq$ 2 kb) or far from ($>$ 2 kb) a known origin (ORI).
