## Additional file 2: Figures for "FORK-seq: replication landscape of the *Saccharomyces cerevisiae* genome by nanopore sequencing"

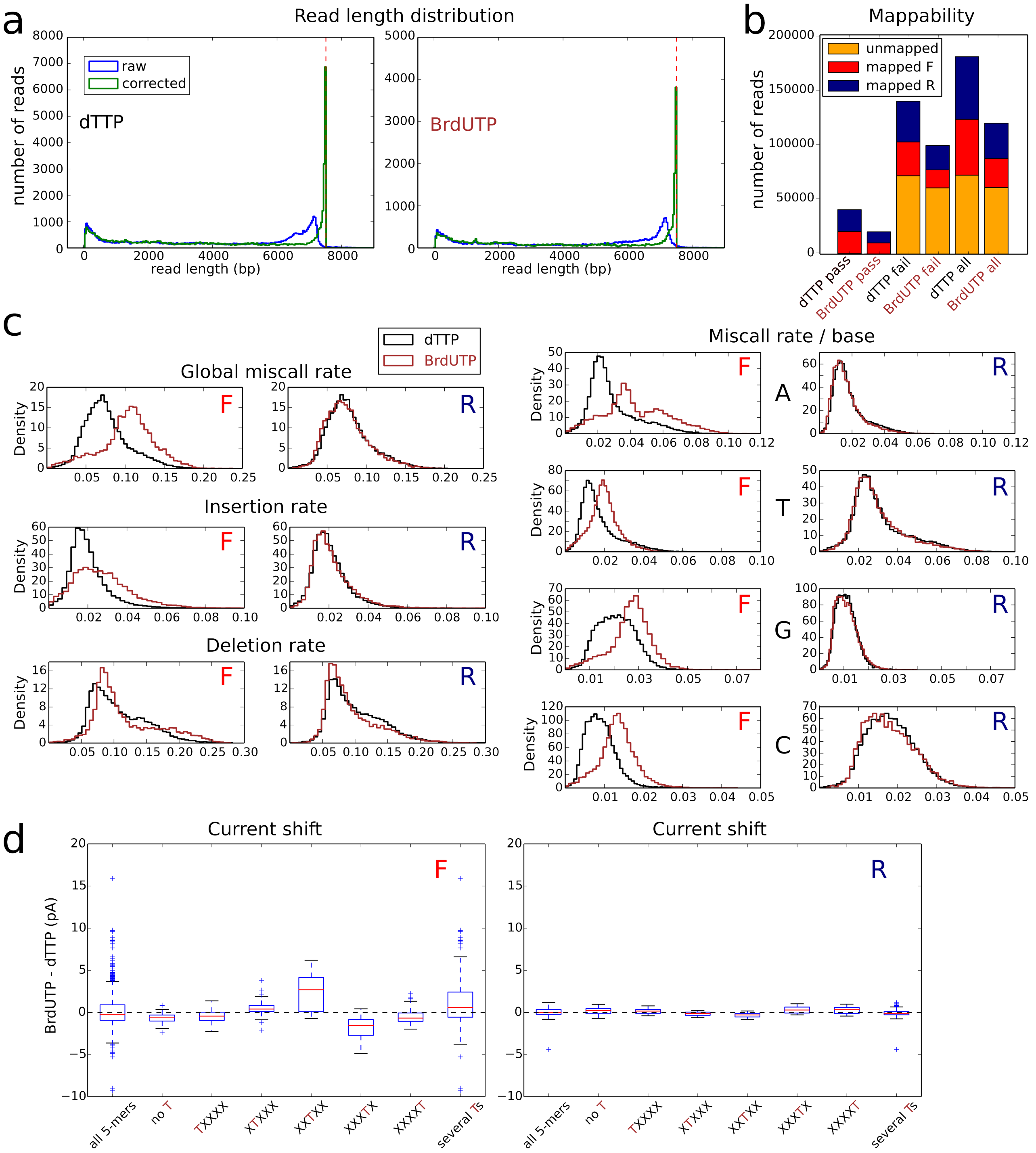


Figure S1: Effect of BrdU on nanopore sequencing. a. Read size histograms, blue : raw length, green : length recalculated after mapping and correction for indels ($L_{corrected} = L_{raw} + deletions - insertions - trimmed$). The dashed red line is the full size (7514 bp) of the plasmid. b. BrdU effect on mappability using BWA MEM after Metrichor basecalling. c. BrdU effect on miscalls, insertions and deletions. Distributions of the miscall rate (global, upper left, per base, right), the insertion rate (middle left) and the deletion rate (bottom left) per mapped read, for the dTTP (black) and BrdUTP (brown) samples. d. Boxplots of the difference of the median of the current distribution between the BrdUTP and the dTTP samples for all 5-mers, 5-mers with no T, 5-mers with a T at the indicated position, and 5-mers with several Ts. For each 5-mer, the median of the current distribution was calculated for the two samples using either the substituted (left, F) or the native (right, R) strand. The difference between the median values of the two samples was then calculated (pA).

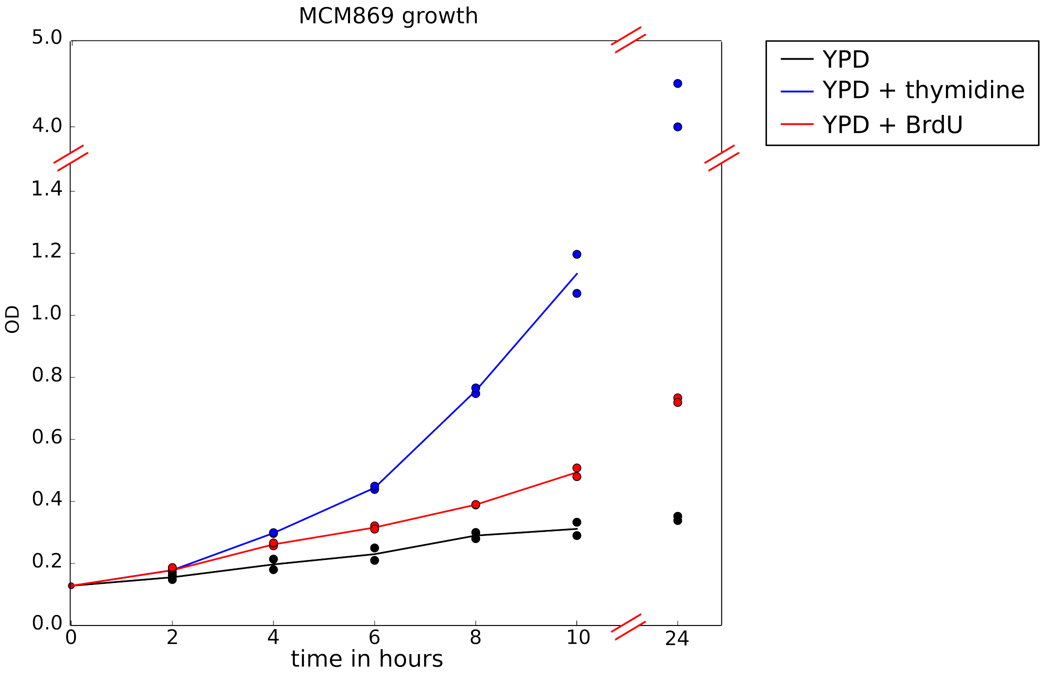


Figure S2: Effect of BrdU on MCM869 growth. MCM869 yeast were grown in YPD for 24 hours after initial dilution, without (black) or with 100 $\mu$M thymidine (blue) or BrdU (red). Cell growth was followed by optical density (OD) measurement at 600 nm. Two biological replicates from the same preculture are shown.

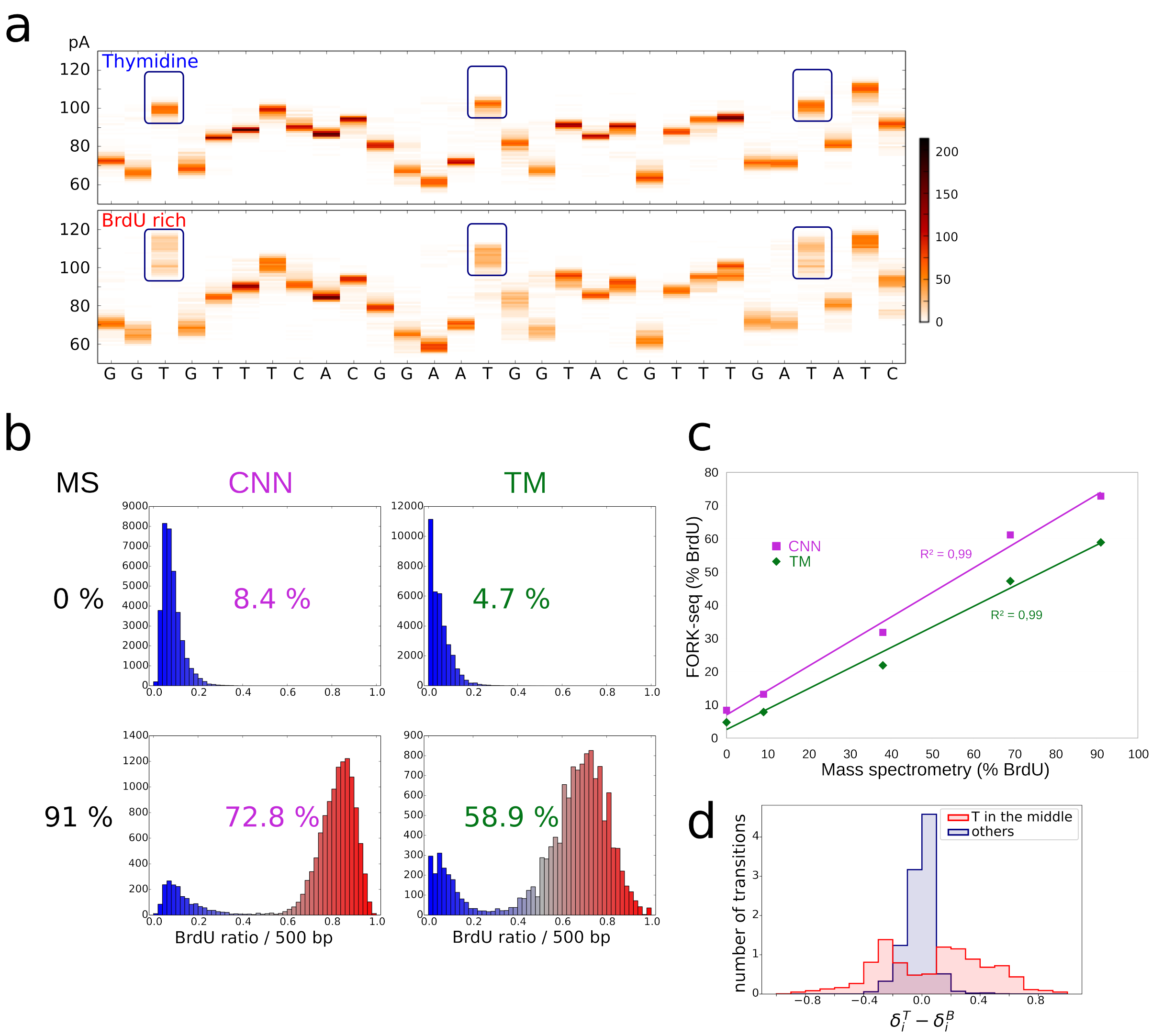


Figure S3: a. Example of a 30 bp sequence (chr12:458000-458029, within rDNA repeats) with the current distribution at each position (400 reads). Blue rectangles highlight some current shifts due to the presence of BrdU. b. Histograms of BrdU proportion averaged over 500 bp for the reads from thymidine control (top) and BrdU-rich (bottom) samples, as determined using CNN (left) and TM (right) detection methods. Overall BrdU proportion of the sample estimated by each method is indicated, as well as the one determined by mass spectrometry measurement (MS, left). c. Overall BrdU content estimated using CNN and TM *versus* mass spectrometry measurements for the 5 samples of given BrdU content (Table T3) after removing reads used in training. d. Distribution of the difference between TM transitions in T and B contexts. For transitions without T in the middle of either starting or ending pentamer (blue), the difference is almost exclusively between -0.4 and 0.4. Transitions with T in one of the pentamers (red) can show significant differences between B and T contexts.

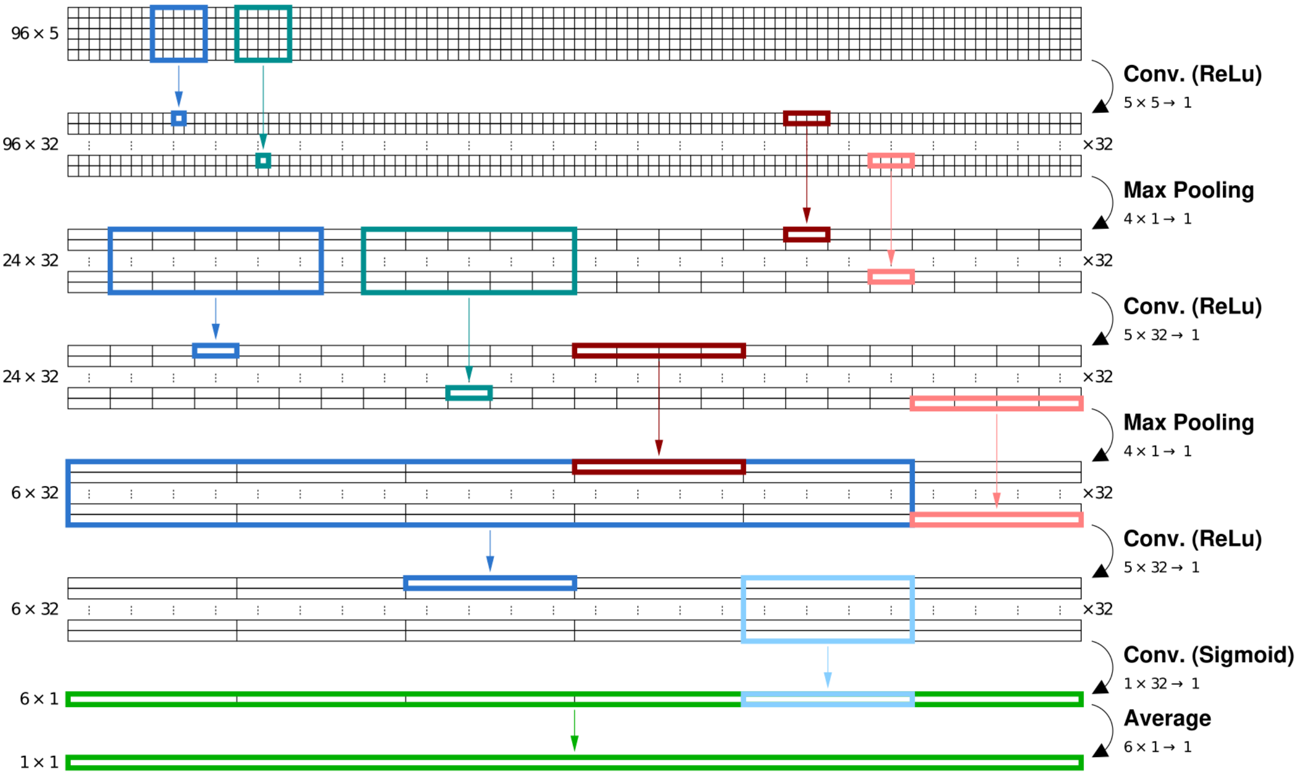


Figure S4: Convolutional neural network architecture. Data representations at each layer are presented from input (top, 96 consecutive current shifts and associated bases) to the output (bottom, proportion of BrdU incorporation over the 96 bp window), with their size specified to the left. Functions operating from one layer to the next are described on the right: 3 convolutional layers (Conv.) with rectified linear (ReLu) or sigmoid activation function alternate with down-sampling operations allowing dimensionality reduction over the sequence ($x$) axis: max pooling and averaging over the $x$ dimension. Color boxes and arrows illustrate the set of cells in layer n-1 used to calculate a cell in layer n. Note, that using 2 max pooling of 4 consecutive values result in the input layer $x$ size being a multiple of $4^2 = 16$.

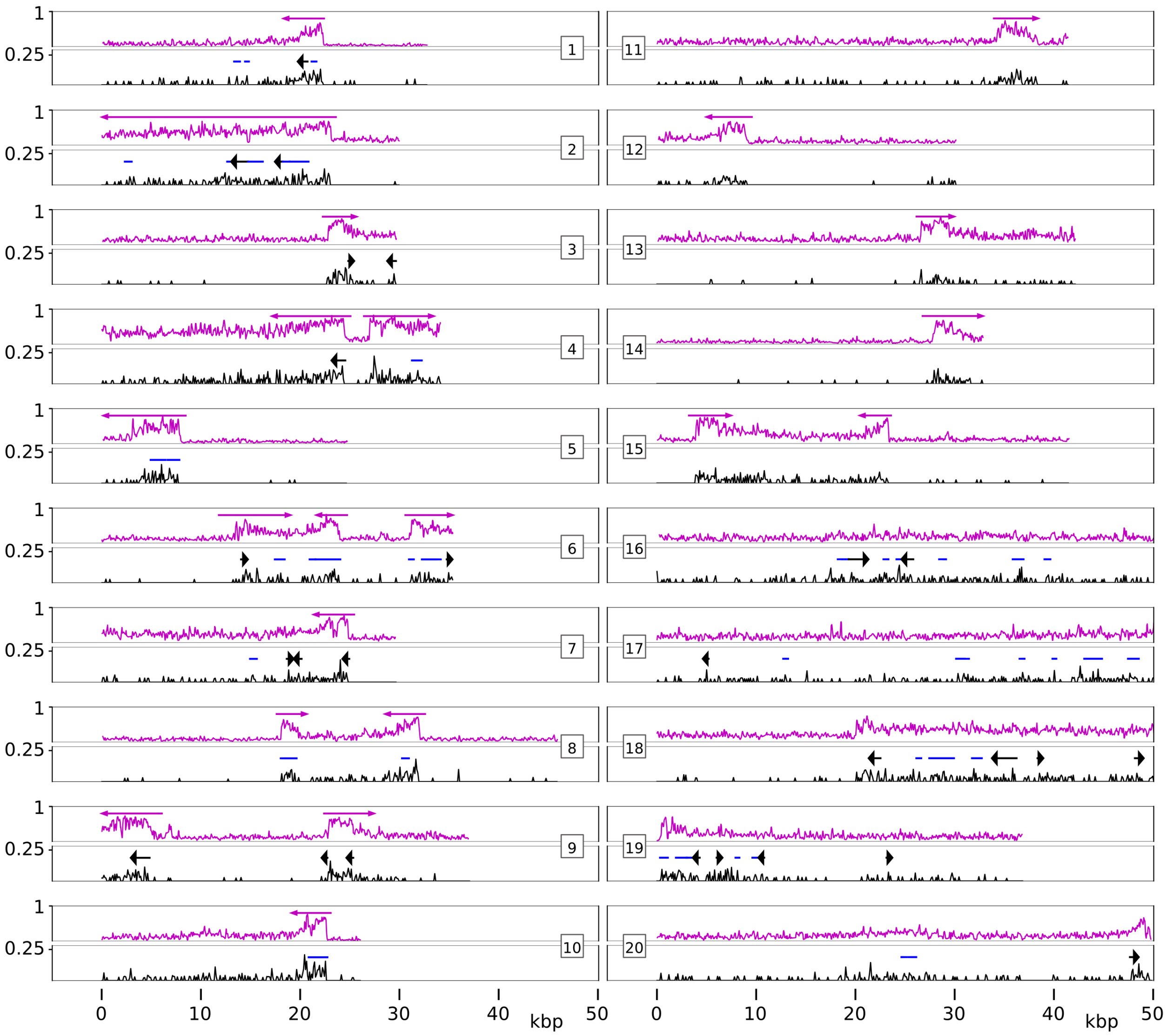


Figure S5: Comparison between RepNano and D-NAscent detection. BrdU content was estimated using either RepNano (CNN, pink) or D-NAscent detect function (black). We then applied our track detection algorithm on both signals. As the signal obtained by D-Nascent is lower, we tried to adapt the detection parameters but we could not obtain a satisfactory detection. Examples of reads showing partial detection (1-10) or no detection (11-15) with D-NAscent of RepNano-detected tracks, and of reads giving false tracks when RepNano didn't (16-20).

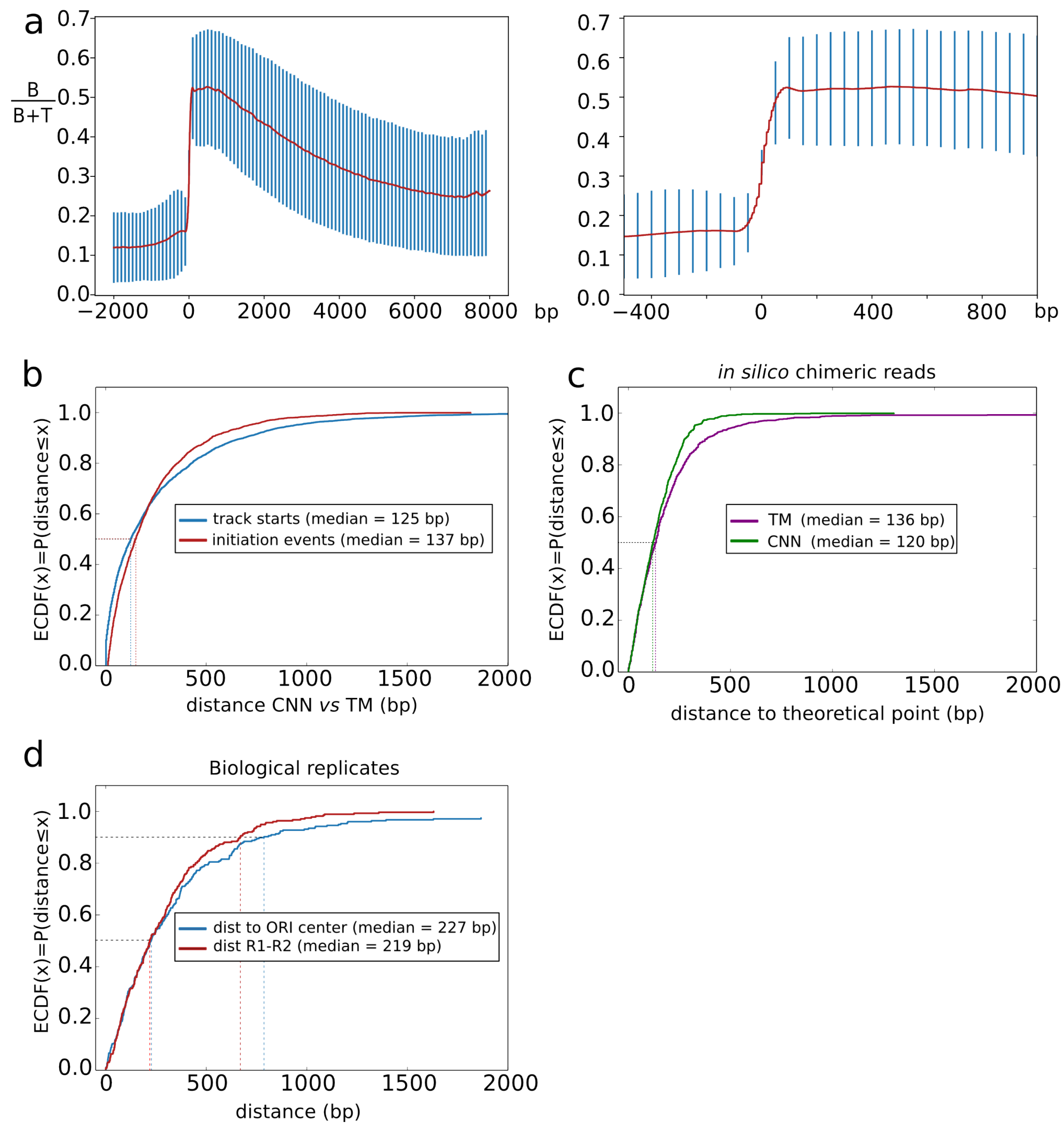


Figure S6: FORK-seq precision. a. Average CNN signal from 7000 replicative tracks. A portion of the left panel is enlarged in the right panel. Raw data were realigned (x=0) at the transition between low ($< 0.3$) and high ($> 0.3$) BrdU level. Blue bars are the standard deviations of the values. The transition from low to high BrdU level occurs within $\sim$ 100 bp. b. Concordance between CNN and TM. Cumulative distribution of distances between track starts (blue) and initiation events (red) obtained by CNN and TM. c. 1000 *in silico* chimeric reads were generated by pasting 2 kb fragments of RepNano outputs (TM or CNN) obtained from two reads from BrdU_R2 sample mapped at the same genomic position but having either low (max B/(B+T) in 500 bp window $< 0.3$) or high (min B/(B+T) in 500 bp window $> 0.7$) BrdU content. This gave us 4 kb fragments with an abrupt transition from low to high BrdU content precisely located in the middle. The fork detection algorithm was then used with the same parameters as for the pulse samples and the distance between the known transition and the detected one was computed. d. Concordance between biological replicates pulse_R1 and pulse_R2. Initiation events were clustered as described in the main text but independently on the two biological replicates. 275 clusters were found common to both replicates. Cumulative distributions of the distances between pulse_R1 cluster median and closest OriDB origin center (blue) and between pulse_R1 and pulse_R2 cluster medians (red) for those shared clusters are shown.

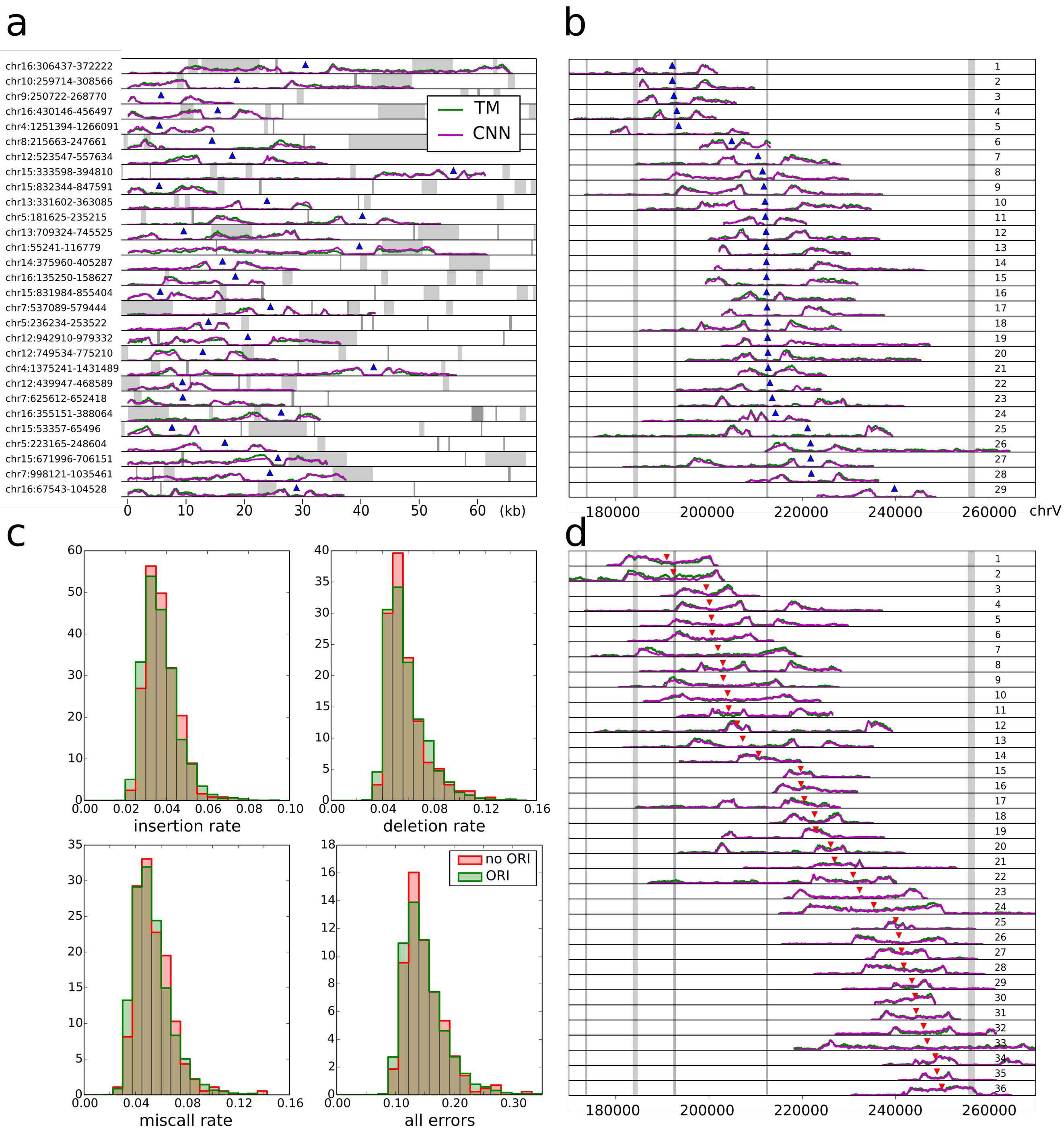


Figure S7: a,b,d. Quantification of the B/(B+T) ratio (B, BrdU, T, thymidine) along nanopore reads using CNN (pink) or TM (green). OriDB origins (confirmed, likely and dubious [?]) are shown as grey vertical bars and detected initiation (resp. termination) events as blue (resp. red) triangles. a. Examples of initiation events detected in the absence of any known origin between the diverging forks. b. Initiation events detected in a 60 kb-region of chromosome V, including a cluster of 4 initiation events (25-28) and two isolated events (6 and 29) not coinciding with a known origin. c. Mapping quality. Normalized histograms of insertion, deletion, miscall and global error rates calculated for all reads containing an initiation, with (green) or without (red) the presence of a known origin between the diverging forks. d. Termination events detected in the same region as in b.

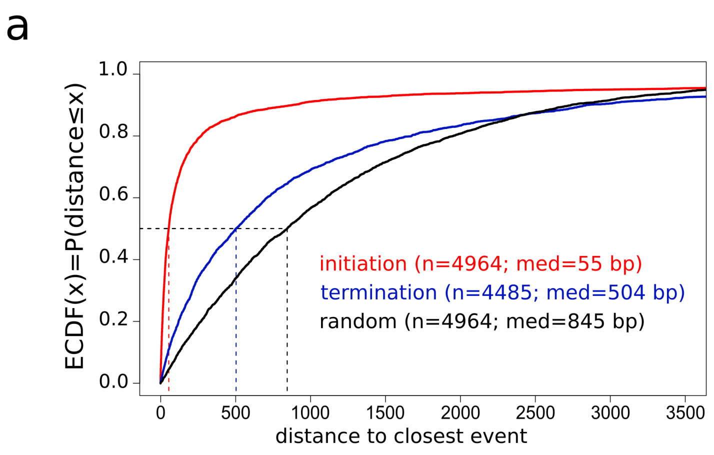


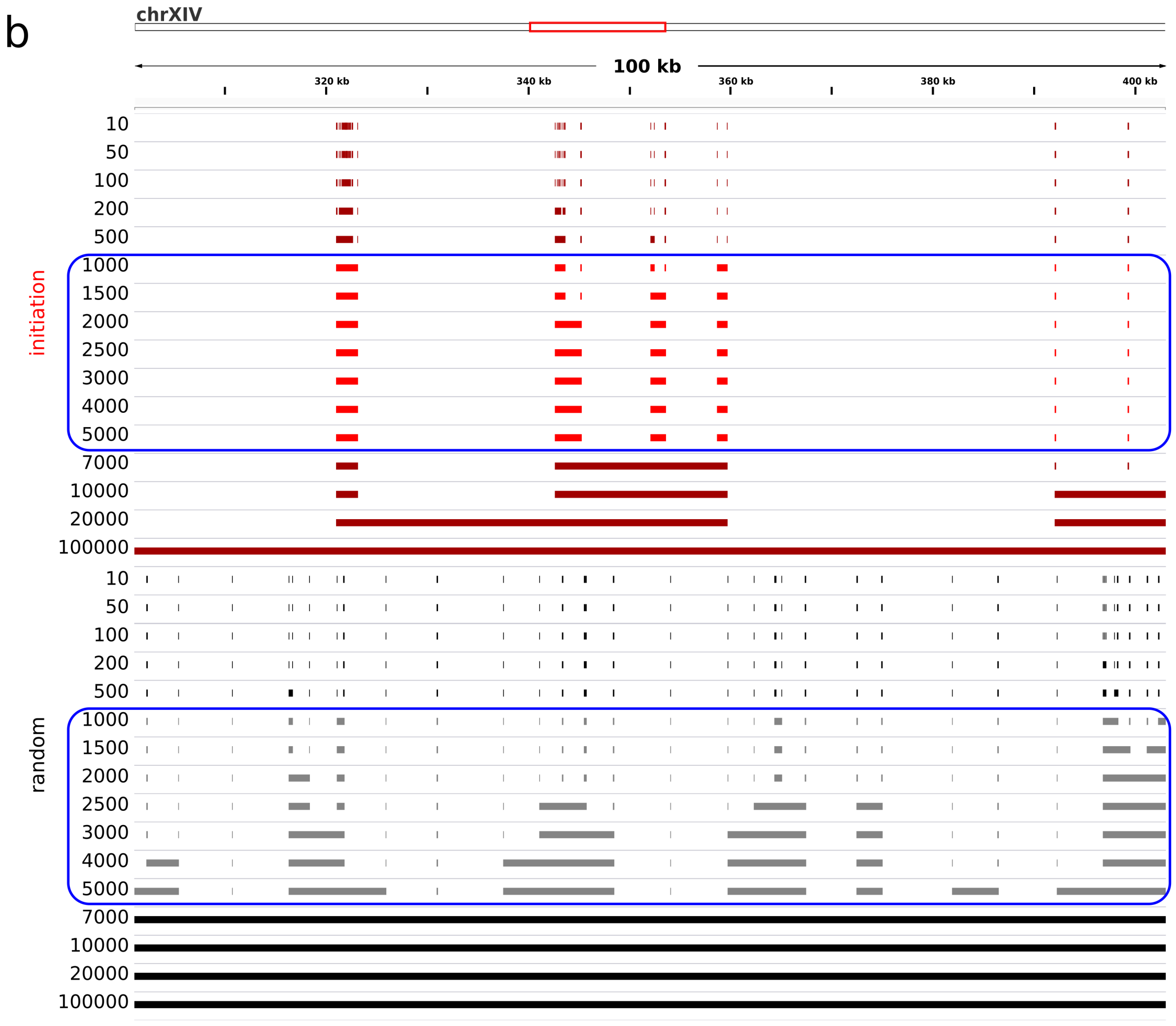


Figure S8: Clustering of initiation and termination events. a. Empirical Cumulative Distribution Function (ECDF) of the distance to the closest neighbour for the indicated objects. Black, random elements, red, initiation events, blue, termination events. b. 100 kb window showing clusters obtained at different mIED (indicated on the left, bp) for initiation events (red) or random elements (black). The blue rectangles highlight mIED values (1 to 5 kb) for which the number of clusters varies less for initiation than for random events (See Fig. 4).

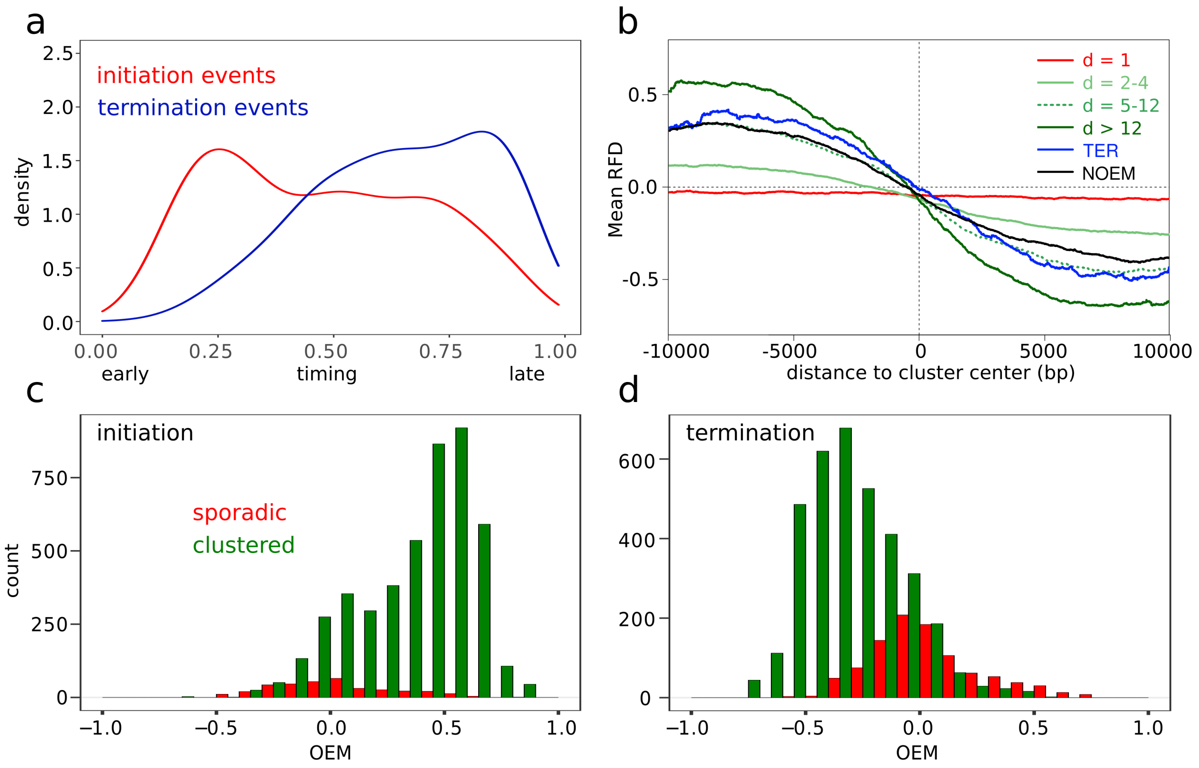


Figure S9: Further characterisation of initiation and termination events in relation to replication timing and fork directionality profiles. a. Density distribution of normalized replication timing (0 = start and 1 = end of S phase) for initiation (red) and termination (blue) events. b. Mean RFD profile (-10kb, +10kb from the median point) of the termination clusters grouped according to their dimension compared to TERs (blue) [**?**] and negative OEM (NOEM, black) segments of the FORK-seq RFD profile. c,d. Distribution of RFD slopes as measured by OEM at sporadic (red) and clustered (green) initiation (c) and termination (d) events.

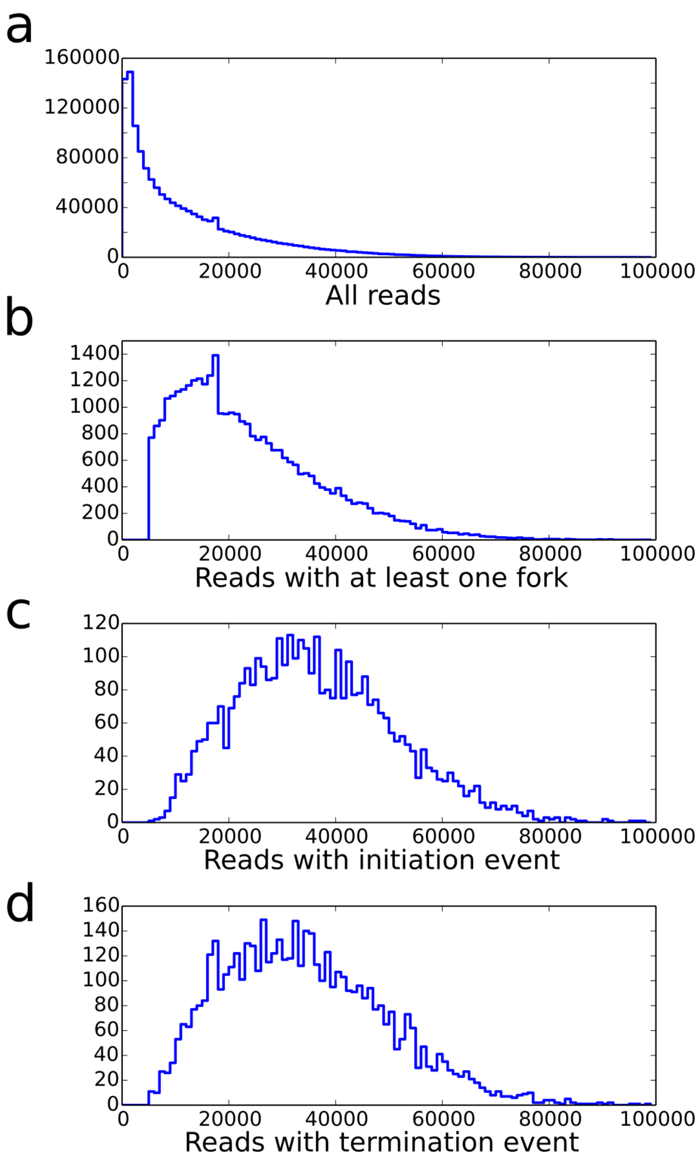


Figure S10: Length distribution of sequenced molecules. a. The length distribution of all reads has a decreasing exponential shape as expected for random DNA breakage and/or random interruption of nanopore translocation. b. The length of reads with one fork is depleted of short molecules as expected from the >5kb threshold size required for BrdU-tract detection. The length of reads with two forks (initiation (c) or termination (d) event) is further shifted to the right as expected from the required presence of at least two BrdU tracts.

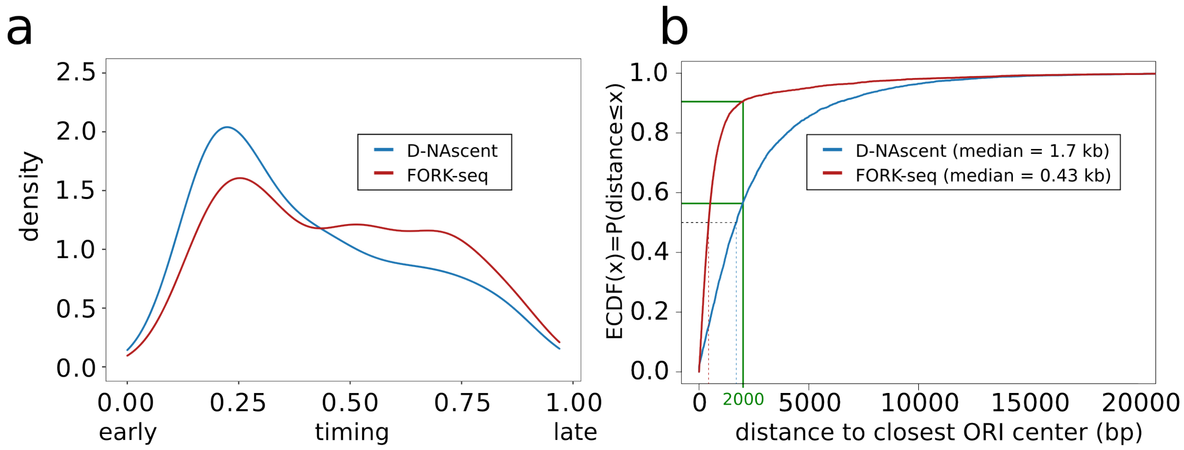


Figure S11: Comparison of D-NAscent and FORK-seq. a. Density distribution of normalized replication timing (0 = start and 1 = end of S phase) for the initiation events detected by FORK-seq (red) and D-NAscent (blue). b. Cumulative distributions of distances between all detected initiation events and known origin centers for D-NAscent [**?**] (n = 6070) and FORK-seq (n = 4964).
